# Gestational diabetes augments group B *Streptococcu*s perinatal infection through disruptions in maternal immunity and the vaginal microbiota

**DOI:** 10.1101/2023.06.23.546252

**Authors:** Vicki Mercado-Evans, Marlyd E. Mejia, Jacob J. Zulk, Samantha Ottinger, Zainab Hameed, Camille Serchejian, Madelynn G. Marunde, Clare M. Robertson, Mallory B. Ballard, Natalia Korotkova, Anthony R. Flores, Kathleen A. Pennington, Kathryn A. Patras

**Affiliations:** Department of Molecular Virology and Microbiology, Baylor College of Medicine, Houston, TX 77030, USA; Medical Scientist Training Program, Baylor College of Medicine, Houston, TX 77030, USA; Department of Microbiology, Immunology and Molecular Genetics, University of Kentucky, Lexington, KY, USA; Department of Molecular and Cellular Biochemistry, University of Kentucky, Lexington, KY, USA; Division of Infectious Diseases, Department of Pediatrics, McGovern Medical School, University of Texas Health Sciences Center at Houston, Houston, TX, USA; Department of Obstetrics and Gynecology, Baylor College of Medicine, Houston, TX 77030, USA; Alkek Center for Metagenomics and Microbiome Research, Baylor College of Medicine, Houston, TX 77030, USA

**Keywords:** Group B *Streptococcus*, *Streptococcus agalactiae*, gestational diabetes, maternal immunity, vaginal colonization, *in utero* dissemination, neonatal outcomes, vaginal microbiome

## Abstract

Group B *Streptococcus* (GBS) is a pervasive perinatal pathogen, yet factors driving GBS dissemination *in utero* are poorly defined. Gestational diabetes mellitus (GDM), a complication marked by dysregulated immunity and maternal microbial dysbiosis, increases risk for GBS perinatal disease. We interrogated host-pathogen dynamics in a novel murine GDM model of GBS colonization and perinatal transmission. GDM mice had greater GBS *in utero* dissemination and subsequently worse neonatal outcomes. Dual-RNA sequencing revealed differential GBS adaptation to the GDM reproductive tract, including a putative glycosyltransferase (*yfhO*), and altered host responses. GDM disruption of immunity included reduced uterine natural killer cell activation, impaired recruitment to placentae, and altered vaginal cytokines. Lastly, we observed distinct vaginal microbial taxa associated with GDM status and GBS invasive disease status. Our translational model of GBS perinatal transmission in GDM hosts recapitulates several clinical aspects and enables discovery of host and bacterial drivers of GBS perinatal disease.

## INTRODUCTION

Group B *Streptococcus* (GBS, *Streptococcus agalactiae*) is a leading agent of neonatal morbidity and mortality and is responsible for ∼150,000 stillbirths or infant deaths annually^1,2^. GBS asymptomatically colonizes the vagina of ∼18% of pregnant women^3^, and subsequently ∼20 million infants are exposed to maternal GBS at, or before, time of delivery^1,2^. About half of infants born to GBS-positive women become colonized and a subset (∼2%) proceed to develop invasive infection^4^. Clinical data supports that GBS can invade the uterus prior to disease onset or stillbirth, even with membranes intact^5–13^, yet the factors driving GBS ascension into the uterus are poorly defined. The current U.S. standard of care, antibiotic prophylaxis during labor to GBS-positive mothers, fails to prevent GBS-associated stillbirths or preterm births, or late onset GBS disease, and exposes ∼1 million U.S. infants to antibiotics each year^14^. This exposure has consequences on the developing infant microbiota including a rise in the number of antimicrobial resistance genes^15,16^. Understanding the biological principles controlling GBS-host dynamics is critical to developing defined, long-lasting preventions for GBS infections in pregnancy and the early neonatal period.

GBS possesses an arsenal of virulence factors (e.g. β-hemolysin/cytolysin, adhesins, hyaluronidase)^17–21^ and environment-sensing two component regulatory systems (e.g. covRS, saeRS, ciaRH, LtdRS)^17,22–24^ that facilitate colonization of the vaginal mucosa and/or invasion of host reproductive tissues. Transcriptomic analyses have revealed that GBS optimizes expression of virulence and metabolic genes to adapt to physiologically relevant stimuli such as pH and glucose *in vitro*^25,26^ and human fluids including blood and amniotic fluid *ex vivo*^27–30^. GBS isolated from the non-pregnant murine vagina differentially expresses adhesion molecules and metabolic genes compared to GBS grown in bacteriologic media and transposon mutant libraries have identified critical GBS factors for murine uterine ascension, yet it is unknown how GBS transcriptionally adapts as it ascends the vaginal tract to the uterus during pregnancy^31,32^.

Maternal metabolic disorders including obesity and gestational *diabetes mellitus* (GDM) are associated with a 1.4 to 3.1-fold increased risk for maternal GBS colonization^33–37^ and a 3 to 5-fold increased risk for GBS maternal and neonatal sepsis^35,36,38^. This clinical evidence implies the maternal diabetic environment is altered in favor of GBS colonization and dissemination, but the factors driving this phenomenon are unknown. Spontaneous hyperglycemia and insulin intolerance are pathophysiological hallmarks of GDM^39^ and are frequently accompanied by maternal systemic immune dysregulation. Most clinical studies focused on maternal peripheral immune profiles and serum biomarkers. Observed differences include increased circulating T helper cell subsets and NK cells in GDM patients with a shift from anti-inflammatory to pro-inflammatory profiles^40–44^ in tandem with increased serum inflammatory mediators and cytokines^45–50^. Although increased placental macrophages and neutrophils are observed in GDM pregnancies^51,52^, GDM perturbations to reproductive immunity are not well-characterized. Furthermore, GDM is accompanied by altered maternal fecal and vaginal microbiome compositions which may further aggravate mucosal dysfunction^53–58^. These clinical findings suggest discordant host-microbe interactions at the mucosa in GDM that may provide an advantage to pathogens, such as GBS, over commensal microbes.

Considering the altered metabolic, immune, and microbial profiles in gestational diabetes compared to non-diabetic pregnancy, we hypothesized that GDM perturbations to host physiology contribute to increased host susceptibility and augmented GBS virulence. To test this hypothesis, we developed a novel mouse model of GBS vaginal colonization and the natural course of maternal-to-offspring transmission in a gestational diabetic host. Using this model, we performed integrative analysis of GBS burden and transmission, host and GBS global transcriptional profiling, reproductive tract cytokine and immune cell profiles, and longitudinal 16S rRNA amplicon sequencing of the vaginal microbiota during pregnancy. Overall, we found that this diet-induced gestational diabetic model mirrors the pathophysiology of GBS disease in GDM, specifically through aberrant maternal immunity, differential GBS metabolic adaptation, and altered composition the vaginal microbiota during pregnancy. These findings advance our current mechanistic viewpoint on host and GBS factors that drive GBS disease in pregnancies impacted by diabetes.

## RESULTS

### A gestational diabetes mouse model of mid-gestational GBS vaginal colonization displays increased maternal carriage and dissemination to fetal tissues

We adapted a previously described diet-induced approach to establish gestational diabetes in C57BL/6J mice^59–61^. Mice are started on either a high-fat high-sucrose (GDM group), or low-fat no-sucrose (pregnant control group) diet one week before mating and maintained on these diets until the experimental endpoint (**Fig. 1A**). In this model, mice develop multiple GDM-like symptoms during pregnancy, including glucose intolerance upon glucose challenge, decreased serum insulin paired with lower total beta cells, insulin resistance, and maternal dyslipidemia by embryonic day 13.5 (E13.5)^59–61^. To model mid-gestational GBS vaginal colonization, mice were vaginally inoculated with GBS strain A909 (serotype Ia) consecutively on E14.5 and E15.5, and maternal and fetal tissues were collected on E17.5 before parturition (potential range of E15.5-17.5 based on mating scheme as described in Methods)(**Fig. 1A**). GDM mice had significantly increased GBS vaginal carriage compared to pregnant controls (**Fig. 1B**), but GBS burdens and rates of ascension into the upper reproductive tract were similar between groups (**Fig. 1B**). To distinguish diet vs. pregnancy-mediated effects, non-pregnant mice on either diet were included in these studies (**Fig. S1A**). Pregnant mice on the HFHS diet (GDM group) had significantly greater vaginal and cervical GBS burden compared to non-pregnant mice on the HFHS diet, with no pregnancy-related differences for mice on the control diet (**Fig. S1B**). These findings suggest that our observations in GDM mice are not solely explained by diet or pregnancy alone, but rather the complex biological interplay of diet and pregnancy in this model.

**Figure 1:**
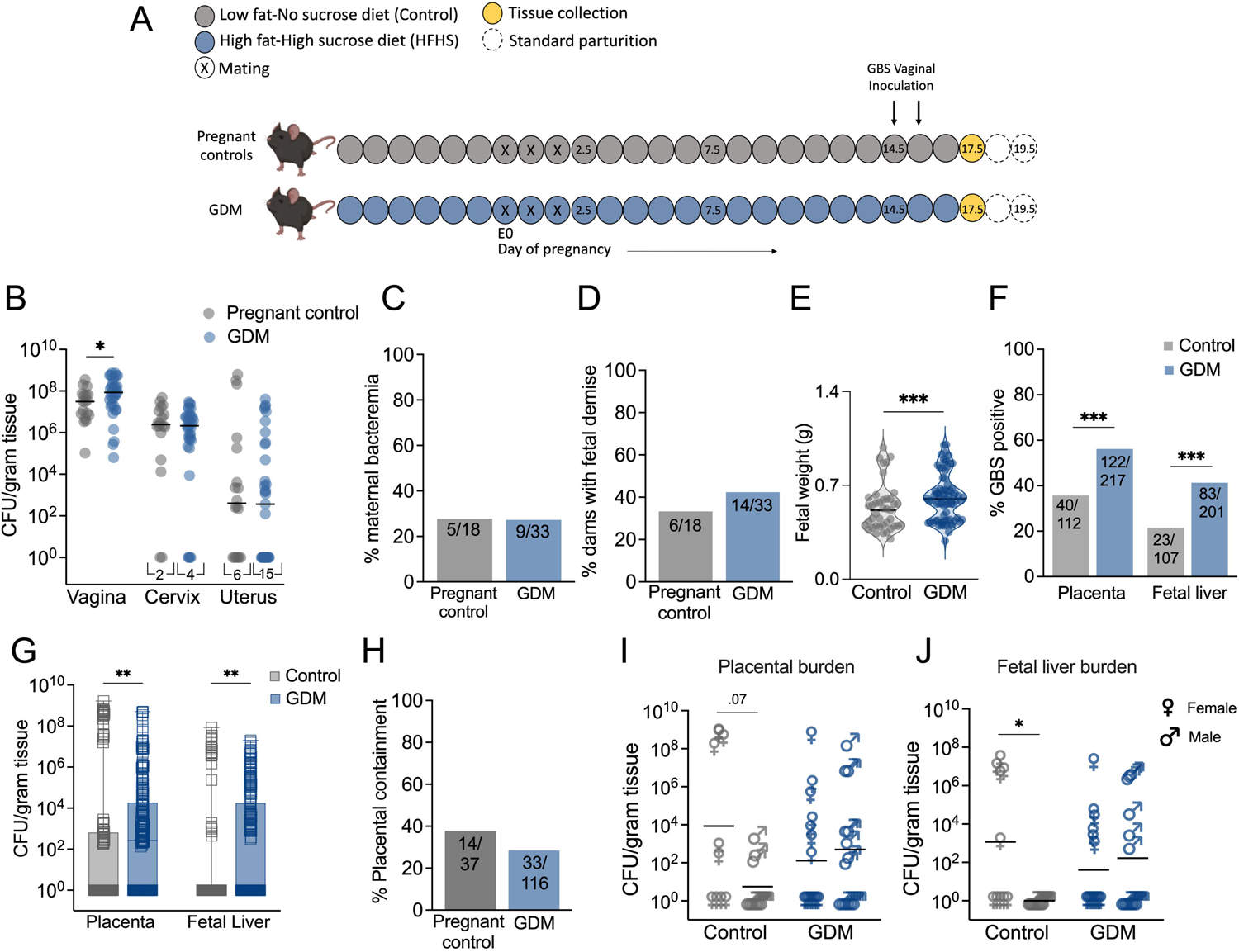
Gestational diabetes enhances *in utero* group B Streptococcal fetal invasion in a murine model of ascending infection. **A)** Experimental timeline for GDM induction via a high-fat high-sucrose (HFHS) diet followed by mid-gestational GBS vaginal colonization, and tissue collection on E17.5. **B)** GBS burden in maternal reproductive tract tissues. Proportion of dams with **C)** bacteremia or **D)** fetal reabsorptions. **E)** Fetal weights from control and GDM dams. **F)** Percentage of placentae and fetal livers that were GBS positive and **G)** corresponding GBS burdens. **H)** Percentage of placental-fetal units that had GBS detected in the placenta with no detection in the corresponding fetal liver. **I**) Placental and **J)** fetal liver burdens stratified by fetal sex for a randomly selected subset. All data represent 4 independent replicates. Points represent individual samples and lines indicate medians (B, E, I-J). Box and whisker plots extend from 25^th^ to 75^th^ percentiles and show all points (G). B-D) *n* = 18 control and 33 GDM dams. E) *n* = 41 control and 75 GDM fetuses. F,H) *n* = 112 control and 217 GDM placentae, 107 control and 201 GDM fetal livers. I-J) *n* = 21 control (10 female, 11 male) and *n* = 30 GDM (16 female, 14 male) paired samples. Data was analyzed by Mann-Whitney t-test (B, E, G, I, J) and Fisher’s exact test (C, D, F, H). *p<0.05, **p<0.01, ***p<0.001.

Incidence of adverse events including maternal bacteremia and fetal resorption (∼30% and ∼40% of pregnancies respectively) were not significantly different between groups (**Fig. 1C,D**). Consistent with observations in uninfected mice^60^, GDM fetuses were significantly larger on E17.5 despite GBS challenge (**Fig. 1E**). Fetuses from GDM dams were significantly more likely to have GBS positive placental and liver tissues (**Fig. 1F**) and had higher GBS burdens in these tissues (**Fig. 1G**) compared to fetuses from control dams. Placental containment, defined as GBS detection in the placenta without translocation to respective fetal liver, was not different between groups (**Fig. 1H**). Disaggregation by fetal sex revealed that females in the control group had significantly greater amount of invasive systemic GBS compared to males determined by liver bacterial burdens, with no sex-specific differences observed in the GDM cohort (**Fig. I-J**).

We also evaluated GBS strain CNCTC10/84 (serotype V), a hypervirulent, hyperhemolytic strain driven by decreased expression of the CovRS (control of virulence) two component system^62^. Similar to A909, CNCTC 10/84 displayed a trend towards higher vaginal GBS burdens in GDM mice (**Fig. S2A**), but no differences in incidence of bacterial uterine ascension nor fetal demise were observed (**Fig. S2B-D**). As with A909, CNCTC 10/84 was significantly more likely to be detected, and at greater amounts, in GDM placentae and fetal livers (**Fig. S2E-F**). CNCTC 10/84 confinement to the placenta was decreased in GDM dams suggesting enhanced invasive potential of this strain (**Fig. S2G**). Given that multiple aspects of GBS infection in the gestational diabetic host were conserved across two GBS strains, A909 was used for the remainder of the study.

### GBS transcriptional adaptation to the uterine environment is impacted by gestational diabetes

To identify GBS genes important for persistence in the gravid uterus and to characterize GBS adaptation to a gestational diabetic host, we performed dual RNA-sequencing of murine vaginal and uterine tissues and GBS harbored in each tissue on E17.5. Principal component analysis revealed that GBS displays tissue-specific transcriptional clustering independent of host diabetic status (**Fig. 2A**). When comparing GBS uterine vs. vaginal transcriptional profiles within pregnant controls and GDM groups, we identified 22 differentially expressed genes (DEGs) contributing to tissue-specific signatures (**Fig. 2B**). Of these 22 DEGs, 13 were shared in both groups and included upregulation of genes responsible for ribose transport and carbohydrate metabolism and downregulation of a transcriptional repressor and several uncharacterized genes (**Fig. 2C-E, Table S1**). Two DEGs were uniquely regulated in GDM mice and seven were uniquely regulated in pregnant controls (**Fig. 2C-E, Table S1**). The two genes uniquely regulated in GDM tissues are yet to be characterized but include a DUF3270 domain-containing protein associated with GBS resistance to bile acid stress^63^ and a YoaK family putative inner membrane protein associated with acid stress in *Escherichia coli*^64,65^. In pregnant controls, uterine GBS significantly upregulated genes associated with sugar transport and cell wall glycosylation, while downregulating genes involved in translation and nucleotide metabolism, compared to vaginal GBS (**Fig. 2C, Table S1**). One upregulated gene, SAK_RS10730 (here named *yfhO*), encodes a predicted YfhO family protein that, in other Gram positive bacteria, is required for glycosylation of critical structural and virulence-contributing cell wall components (lipoteichoic or wall teichoic acid)^66–68^. AlphaFold2^69,70^ modeling predicted 14 transmembrane helical domains suggesting YfhO is an integral membrane protein (**Fig. 2E**). STRING analysis^71^ of predicted YfhO protein-protein interactions suggests a potential involvement in enzymatic reactions (glycosyl and phosphoryl transfer) and antibiotic resistance mechanisms (PBP2B and ABC transporter)^72^ (**Fig. 2F**). We assessed the role of YfhO in ascending uterine infection using an isogenic allelic exchange mutant expressing a kanamycin resistance cassette in place of *yfhO* (Δ*yfhO*). Compared to parental A909, Δ*yfhO* showed decreased GBS burdens and rate of ascension to the uterus in pregnant control and GDM mice, without deficits in vaginal colonization or ascension to the cervix (**Fig. 2H,I**). This attenuation was specific to the gravid uterus; Δ*yfhO* had no deficits in uterine ascension in non-pregnant mice (data not shown). Additionally, Δ*yfhO* maintained the capacity to invade GDM placental tissues, but not control placentae (**Fig. 2J**), and was not detected in fetal livers from either group (data not shown). To test whether GDM mice were more broadly susceptible to GBS colonization and fetal dissemination, we also assessed the role of the well-characterized β-hemolysin/cytolysin using an isogenic mutant of *cylE*, a gene required for β-H/C production^73^. A909 Δ*cylE*, previously shown to have attenuated fetal dissemination^18^ and vaginal colonization in non-pregnant mice^17^, was equally attenuated in control and GDM mice (**Fig. 2H,J**).

**Figure 2:**
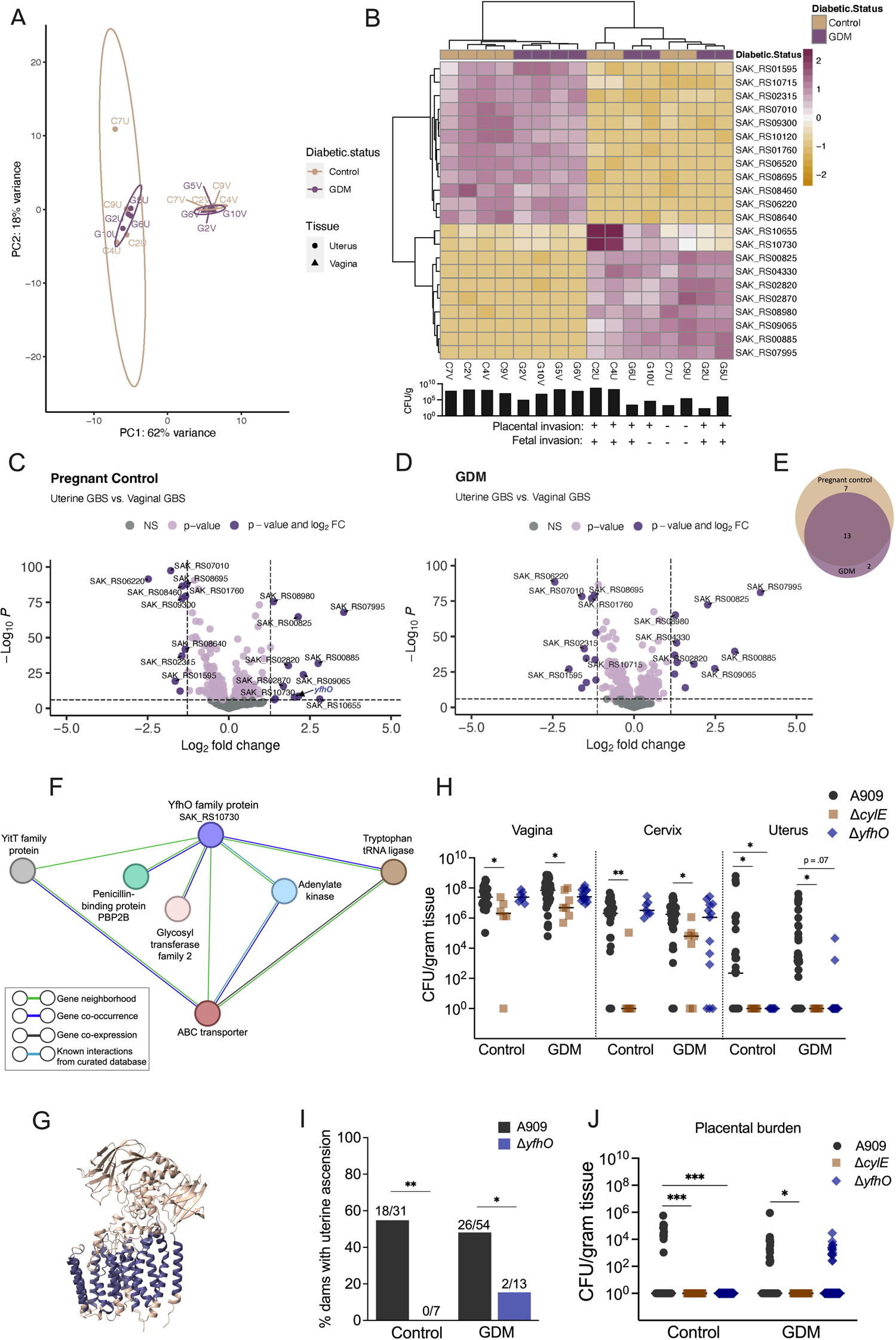
GBS transcriptional profiling in control and gestational diabetic pregnancy identifies genes important for ascension to the gravid uterus. **A)** Principal component analysis of GBS transcriptional profiles from vaginal (V) or uterine tissue (U) of pregnant control (C) or GDM (G) mice on E17.5 (*n* = 4/group, one independent experiment). Each point represents a sample from an individual mouse, with vaginal-uterine pairs indicated by numeric label. **B)** Heatmap of the 22 genes differentially expressed in GBS from uterine tissue compared to GBS from vaginal tissue of pregnant controls and GDM mice. Tissue burdens and presence or absence of GBS placental and fetal dissemination are indicated below the heatmap. Volcano plot of differentially expressed genes (DEGs) in uterine GBS vs. vaginal GBS in **C)** pregnant controls or **D)** GDM mice, and a **E)** Venn diagram showing the corresponding proportion of GBS DEGs unique vs. shared between comparisons. See also **Table S1**. **F)** Predicted AlphaFold2 structural modeling of YfhO (SAK_RS10730) protein structure. **G)** Predicted functional protein network of SAK_RS10730 determined by STRING analysis. **H)** E17.5 GBS burdens in the reproductive tract of mice challenged with WT A909, Δ*cylE*, or Δ*yfhO*. **I)** Proportion of dams with uterine ascension of WT A909 or Δ*yfhO*. **J)** Placental GBS burden in control and GDM groups (*n* = 44-116/group). H-J) Data for WT A909 are aggregated from all experiments, of which 5 pregnant controls and 10 GDM are from concurrent GBS mutant experiments (*n* = 31 pregnant controls, *n* = 54 GDM), mice challenged with Δ*cylE* (*n* = 6 pregnant controls, *n* = 7 GDM) are from 2 independent experiments, and mice challenged with Δ*yfhO* (*n* = 7 pregnant controls, *n* = 13 GDM) are from 5 independent experiments. Points represent individual samples and lines indicate medians (H,J). DEGs were identified via generalized linear model, Log_2_ fold change >1 and FDR adjusted p-value <0.05 (B-E). Data were analyzed by Kruskal-Wallis followed by Dunn’s multiple comparisons test (H, J) and two-sided Fisher’s exact test (I). *p<0.05, **p<0.01, ***p<0.001.

### Gestational diabetes alters the maternal vaginal and uterine transcriptional landscape and cytokine responses during GBS infection

Host transcriptional profiles also clustered in a tissue-specific manner independent of diabetic status (**Fig. 3A**). When comparing the vaginal transcriptome of GDM mice to pregnant controls, we identified four genes that were significantly downregulated in the diabetic group including *Chil4*, a positive regulator of chemokine production and Class Ib MHC antigen *H2-Q6*, associated with antigen processing and T cell-mediated cytotoxicity (**Fig. 3B, Table S2**). Significantly upregulated pathways involved DNA repair, ECM degradation, IL-17, and TLR signaling suggesting greater GBS-associated vaginal tissue damage and inflammation in GDM hosts (**Fig. S1C**). When comparing the uterine transcriptome of GDM mice to pregnant controls, GDM uterine tissue displayed downregulation of inflammation, wound response, and fertility genes compared to pregnant controls (**Fig. 3C, Table S2**). Significantly downregulated pathways in the GDM uterus included several immune responses (neutrophil recruitment and degranulation, and TLR, IFN1, and TGFB signaling) and significantly upregulated pathways involved DNA damage, senescence, and hypoxia responses (**Fig. 3D**). Together, transcriptional profiles suggested altered GBS-associated immune responses in GDM hosts. Indeed, a multiplex cytokine assay showed that GDM mice had a dysregulated vaginal cytokine response to GBS characterized by significantly greater levels of G-CSF and KC and lower IL-2 compared to pregnant controls (**Fig. 3E,F, Fig. S3**). Compared to respective mock-infected dams, GBS inoculation elevated vaginal KC, IL-1β, and IL-12p70 in both GDM and control dams. Additionally, GDM mice had significant induction of IL-1α and G-CSF, whereas pregnant controls significantly induced IL-2, compared to mock-infected counterparts. Uterine IL-1α was uniquely induced in GBS-infected GDM dams compared to mock-infected GDM dams, and there were no cytokine differences detected in placental tissues (**Fig. 3E, Fig. S3**).

**Figure 3:**
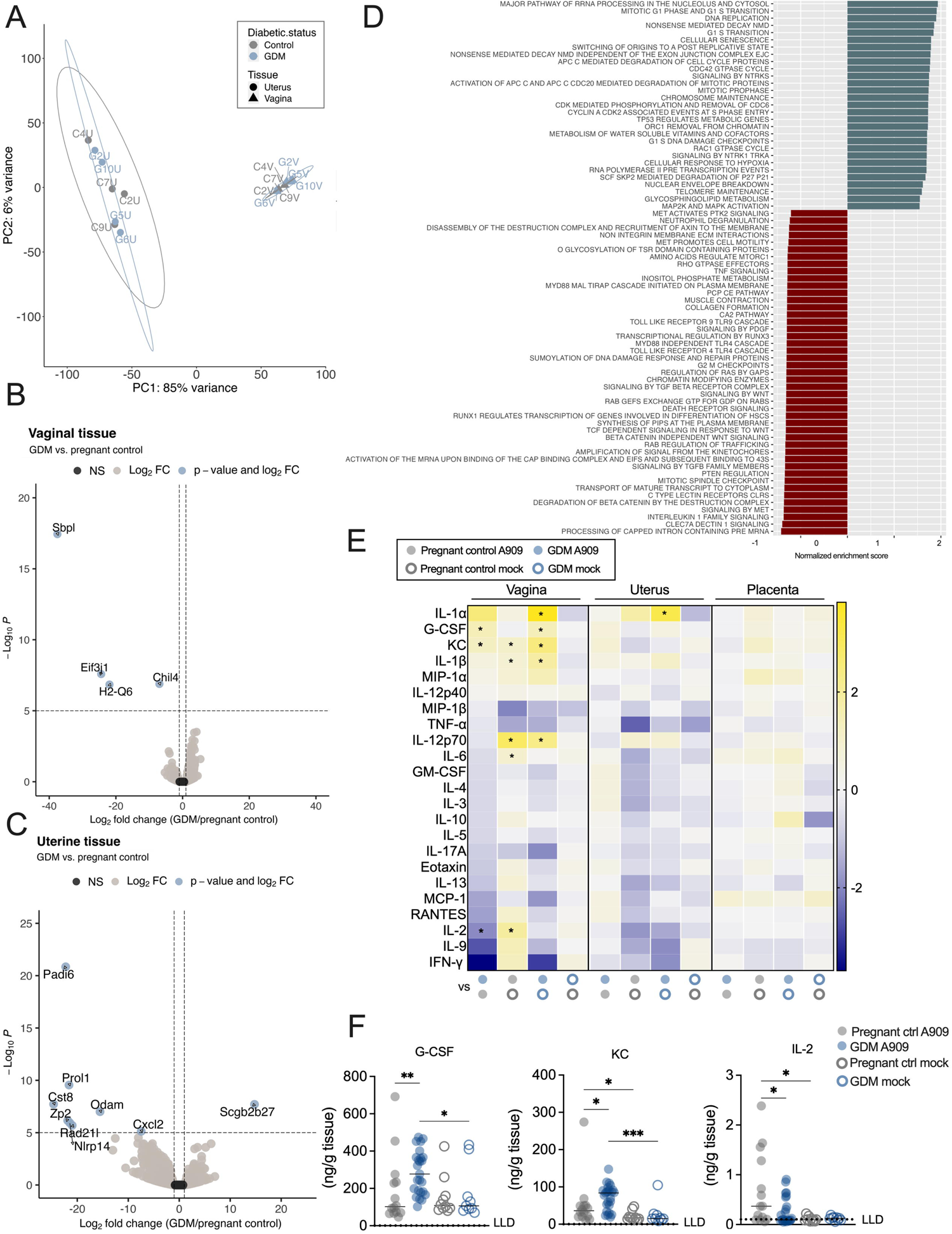
The reproductive transcriptional landscape and cytokine response are altered in gestational diabetic mice during GBS challenge. **A)** Principal component analysis of host transcriptional profiles from vaginal (V) or uterine tissue (U) of pregnant control (C) or GDM (G) mice on E17.5 (*n* = 4/group, one independent experiment). Each point represents a tissue from an individual mouse, with vaginal-uterine pairs indicated by numeric label. Volcano plot of differentially expressed genes (DEGs) in **B)** vaginal tissue and **C)** uterine tissue from GDM mice vs. pregnant controls. **D)** Gene set enrichment analysis of uterine tissue in gestational diabetic mice vs. pregnant controls**. E)** Heatmap of 23 cytokines in vaginal, uterine and placental tissues on E17.5 from pregnant controls and GDM mice that were inoculated with A909 or mock-infected. **F)** Vaginal cytokines that were significantly different between infected GDM and controls. Data (E-F) are from 3 independent experiments with *n* = 14 infected controls, 25 infected GDM, 10 mock-infected controls and 9 mock-infected GDM mice. Points represent individual samples and lines indicate medians (F). DEGs were identified via generalized linear model, Log_2_ fold change >1 and FDR adjusted p-value<0.05 (B-C). fGSEA was performed with a gene set minimum of 15, a gene set maximum of 500, 10,000 permutations, and the Reactome gene set collections from the Molecular Signatures Database. Cytokine data were analyzed by Kruskal-Wallis test followed by a two-stage linear step-up procedure of Benjamini, Krieger and Yekutieli to correct for multiple comparisons by controlling the false discovery rate (<0.05). *p<0.05, **p< 0.01, ***p<0.001.

### Immune cell populations are altered in gestational diabetic mice following GBS inoculation

Next, we evaluated immune cell infiltration in E17.5 vaginal, uterine and placental tissues by flow cytometry using a 19-marker antibody cocktail to distinguish lymphoid and myeloid lineage subsets (**Fig. 4A**). There were no differences in the composite CD45+ fraction between GDM and control samples across tissue types (**Fig. 4B**). Additionally, no differences between GDM and control maternal splenic immune populations were detected (**Fig. S4A**). GDM mice displayed several significantly altered immune cell subsets proportions compared to controls including increased B cells in vaginal tissues (**Fig. 4C**), and decreased NK and regulatory T cells in uterine tissues (**Fig. 4D**), with no differences in total immune cell counts in either tissue (**Fig. 4H, Fig. S4B-E**). Uterine NK (uNKs) cells, defined as uterine NK1.1^+^ CD69^+^ cells^74^, were enriched in uteri of both groups (**Fig. 4E**), but GDM mice displayed a significantly decreased proportion of uNKs positive for activation marker CD25 (**Fig. 4F**). While proportions of placental immune cells were similar between groups, total cell counts revealed that GDM placentae had significantly fewer CD45^+^ cells, and several immune subpopulations including neutrophils and NKs (**Fig. 4G-H, Fig. S4B-C,F**). Given evidence of dysregulated neutrophil transcriptional responses and neutrophil-recruiting chemokines (**Fig. 3D-F**) and reduced NK activation (**Fig. 4D,F**) in GBS-challenged GDM mice, we further probed their involvement in ascending fetal infection. Neutrophils or NK cells were depleted with α-Ly6G or α-NK1.1 antibodies respectively with one dose given before and an additional dose given during GBS challenge (**Fig. 5A**). Specific local depletion of NK cells (**Fig. 5B,C**) or neutrophils (**Fig. 5B,D**), was confirmed by flow cytometry of uterine tissues (**Fig. S4G**). NK depletion did not impact maternal tissue burdens in GDM or pregnant controls (**Fig. 5E-G**). Neutrophil depletion led to greater GBS load in vaginal tissues of pregnant controls (**Fig. 5E**), with no effect on cervical or uterine burdens in both groups (**Fig. 5F,G**). In GDM mice, NK depletion had protective effects against fetal infection shown by significantly fewer GBS-positive fetoplacental tissues (**Fig. 5H-J**) and decreased bacterial load in fetal livers (**Fig. 5K**). Pregnant controls displayed reduced GBS invasion of fetoplacental tissues overall compared to the GDM group as in **Fig. 1F,G**, with NK depletion significantly reducing the incidence of fetal liver GBS invasion (**Fig. 5J**). In contrast, neutrophil depletion was inconsequential for fetal dissemination in the GDM group, but in pregnant controls, led to increased frequency of placental GBS positivity (**Fig. 5H-K**). Additionally, preterm birth (delivery of pups at or before E17.5) was observed in 2/5 GDM dams treated with anti-Ly6G but this was not observed in the corresponding pregnant controls (0/7 dams). Collectively, these findings suggest distinct roles for NKs and neutrophils in controlling GBS infection at the maternal-fetal interface in healthy pregnancy and in susceptible hosts.

**Figure 4:**
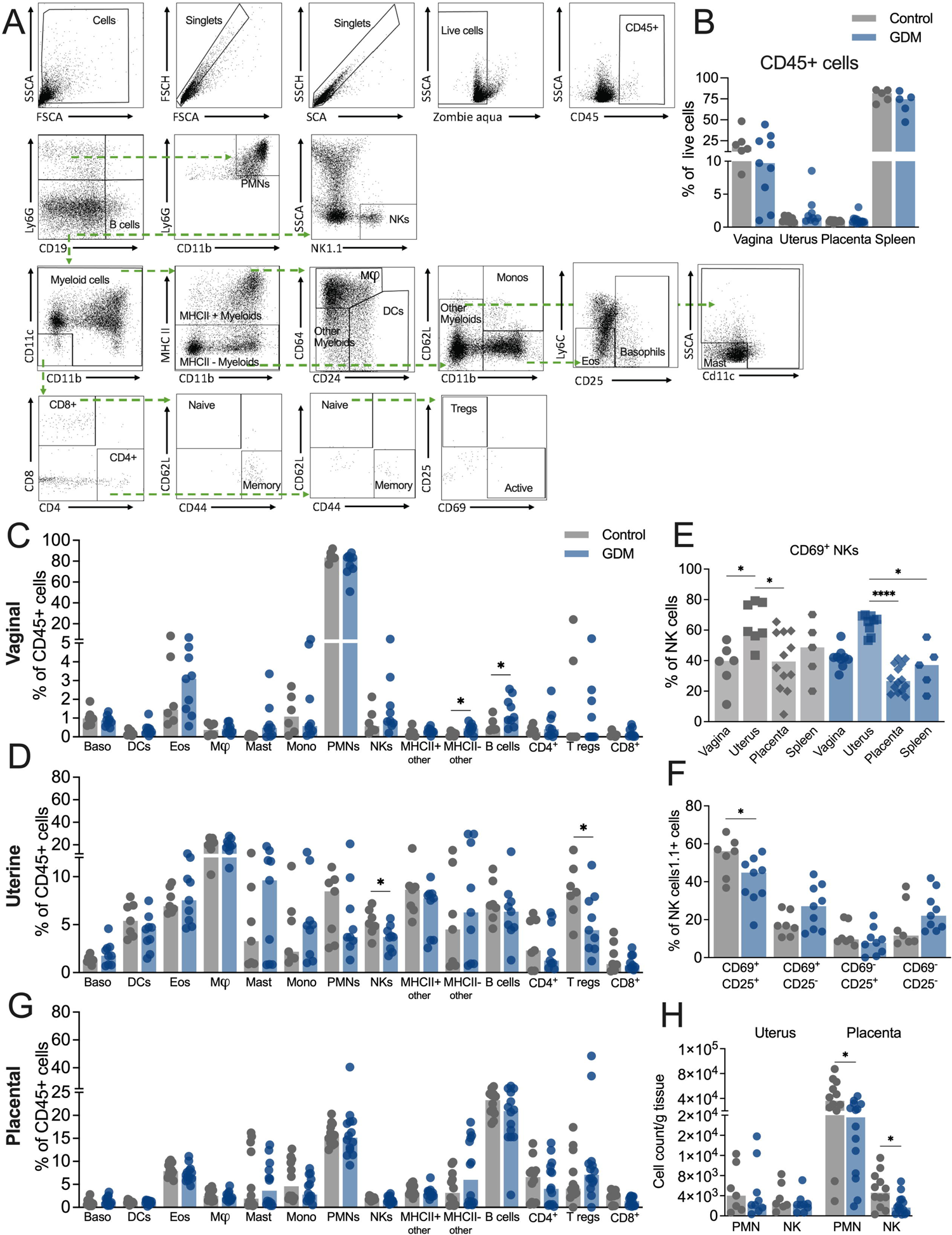
Immune cell recruitment is dysregulated in gestational diabetic mice. **A)** Gating strategy for assessing recruitment of basophils (Baso), dendritic cells (DCs), eosinophils (Eos), macrophages (M_), mast cells (Mast), monocytes (Mono), neutrophils (PMNs), NK cells (NKs), B cells, CD4^+^ T cells, CD4^+^ regulatory T cells (T regs) and CD8^+^ T cells by flow cytometry. **B)** Frequency of CD45+ cells. Immune cell frequencies in **C)** vaginal and **D)** uterine tissues in GBS-infected dams. **E)** Frequencies of CD69+ NK cells across tissues. **F)** Uterine NK cell proportions stratified by CD69 and CD25 expression. **G)** Immune cell frequencies in placentae collected from GBS-infected dams. **H)** Total cell counts of neutrophil and NK cells from placentae. Data (B-H) are from 3 independent experiments with each point representing an individual mouse sample (*n* =7 pregnant controls and 9 GDM, with 1-2 placentae per dam for a total of *n* = 12 control placentae and 14 GDM placentae). Data were anaylzed by Mann-Whitney t-tests per tissue. *p<0.05, ****p<0.0001.

**Figure 5:**
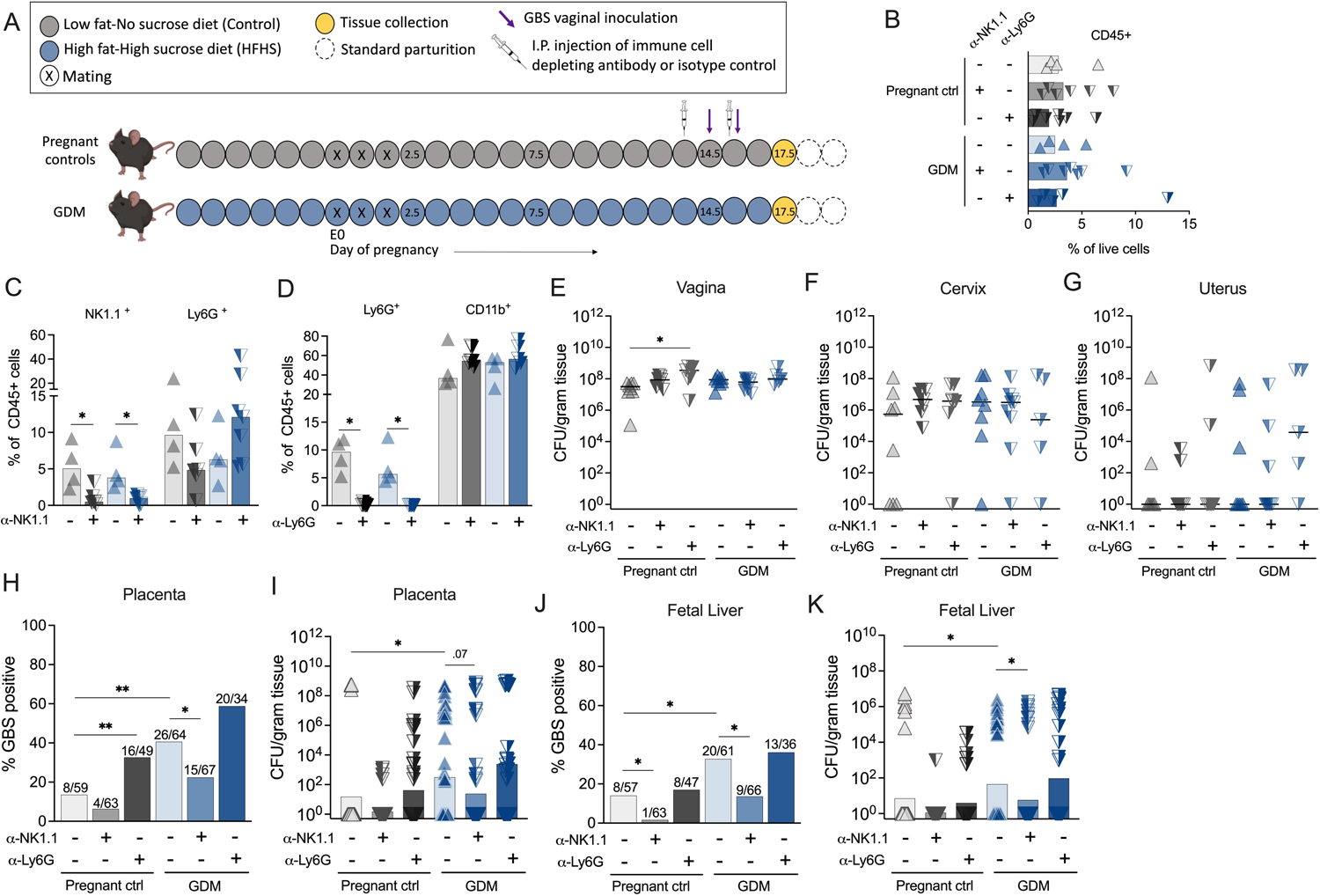
NK and neutrophil depletion differentially impact ascending GBS infection. **A)** Schematic of experimental timeline for immune cell depletion and subsequent GBS challenge in GDM and pregnant control (ctrl) mice. **B)** Frequencies of uterine CD45+ cells in NK-depleted (α-NK1.1), neutrophil-depleted (α-Ly6G), and isotype-treated mice. **C)** Frequencies of uterine NK1.1+ and Ly6G+ cells in isotype-treated or NK-depleted mice. **D)** Frequencies of uterine Ly6G+ and CD11b+ cells in isotype-treated or neutrophil-depleted mice. **E-G)** GBS burdens of maternal reproductive tract tissues with (α-NK1.1 or α-Ly6G) or without (isotype control) immune cell depletion in GDM and pregnant control mice. **H)** GBS positive proportions and **I)** burdens of placentae with or without immune cell depletion in GDM and pregnant control mice. **J)** GBS positive proportions and **K)** burdens of fetal livers with or without immune cell depletion in GDM and pregnant control mice. Data for isotype treated mice are pooled from 7 independent experiments; 4 NK depletion experiments and 3 neutrophil depletion experiments. For NK depletion experiments, *n* = 7 pregnant controls and 9 GDM mice that were NK-depleted, *n* = 7 pregnant controls and 6 GDM mice that were isotype-treated. For neutrophil depletion experiments, *n* = 7 pregnant controls and 5 GDM mice that were neutrophil-depleted, *n* = 7 pregnant controls and 6 GDM mice that were isotype-treated. Experimental numbers for placenta-fetal pairs in each group are given as denominators in H and J. Data were analyzed by multiple Mann-Whitney t-tests with correction for multiple comparisons (C,D), by Kruskal-Wallis test with a post-hoc Dunn’s multiple comparisons test (E-G,I,K), or by Fisher’s exact test (H,J). *p<0.05, **p<0.01.

### Gestational diabetes augments adverse neonatal outcomes associated with GBS

Previously described mouse models of intra-vaginal GBS inoculation and subsequent transmission to vaginally-delivered neonates have shown adverse outcomes such as decreased survival, stunted weight, and systemic infection^75,76^. To examine the effect of GDM on offspring outcomes, we monitored neonatal outcomes born to GBS-inoculated dams through postnatal day 7. Neonates born to GDM dams had significantly worse survival and stunted weight in the first week of life compared to neonates from control dams (**Fig. 6A,B**). To track GBS transmission and systemic disease, we measured GBS burden in neonatal intestines (indicating colonization) and livers (indicating invasive disease) upon death or on postnatal d7 in surviving pups. We found no differences in intestinal GBS colonization or systemic infection between GDM and control neonates regardless of time of collection (**Fig. 6C,D**). There were also no differences in hours to delivery following first GBS inoculation (**Fig. 6E**). A stark disparity in female pup survivorship likely contributed to observed differences between GDM and control groups (**Fig. 6F**). While there were no sex-dependent differences in d7 pup weights (**Fig. 6G**), GDM males had significantly greater liver burden on d7 compared to GDM females, with no sex-effects observed in controls (**Fig. 6H**). It is possible that the observed sex differences in the GDM cohort are related, and that female pups with greater GBS invasive disease succumbed to GBS prior to d7 and thus were not accounted for in the d7 liver samples; however, this speculation cannot be confirmed by the current experimental design. Together, these data suggest that gestational diabetic pregnancies exhibit increased susceptibility of GBS dissemination to the fetal compartment with subsequent worse neonatal survival and growth.

**Figure 6:**
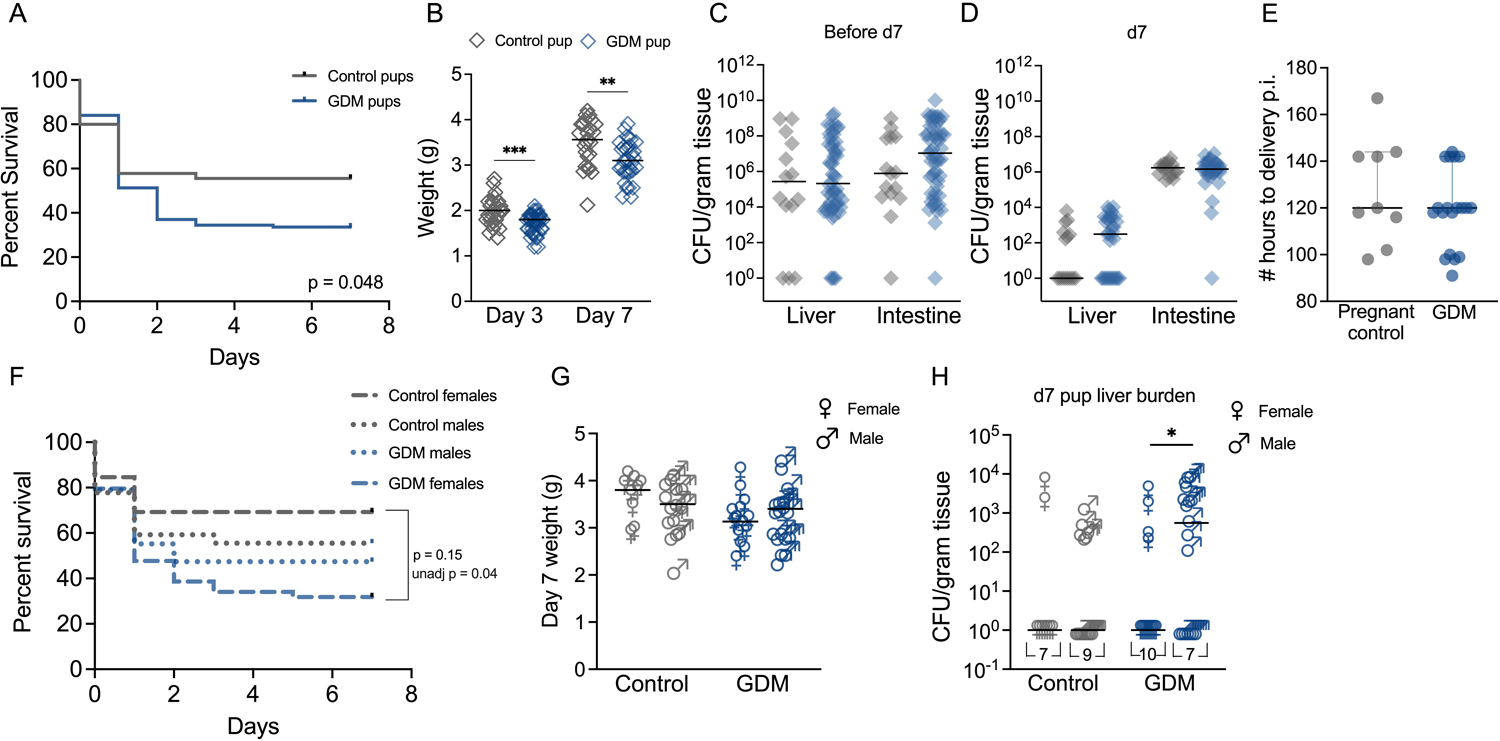
Gestational diabetes worsens neonatal outcomes associated with GBS infection. **A)** Survival of neonates from GDM or control dams that were inoculated with GBS on E14.5 and E15.5. *n* = 45 offspring from 9 control dams and 119 offspring from 19 GDM dams. All remaining neonates were sacrificed on d7 for burden quantification. **B)** Neonatal weights on days 3 and 7 of life, *n* = 25 control pups and 35-42 GDM pups. GBS burden in pup livers and intestines **C)** near time of death before d7, or **D)** on d7 (*n* = 15-42/group). **E)** Time to delivery following initial GBS inoculation. **F)** Pup survival, **G)** d7 pup weights, and **H)** pup GBS liver burdens stratified by sex. All data are from 4 independent experiments. Points represent individual samples and lines indicate medians. Data were analyzed via Kaplan-Meier survival analysis with Holm-Šídák correction for multiple comparisons (A, F) and Mann-Whitney t-test (B-E, G-H). *p<0.05, **p<0.01, ***p<0.001.

### Gestational diabetes impairs normal progression of the vaginal microbiota in pregnancy and specific taxa are associated with GDM and GBS ascension

The vaginal microbiota composition is associated with pregnancy complications and birth outcomes across patient demographics and geographic regions^77–81^ and several clinical studies have demonstrated that women with GDM exhibit distinct distribution of vaginal taxa compared to healthy controls^53,54,82–84^. By collecting vaginal swabs every 3 days from diet introduction through E14 of pregnancy, we longitudinally characterized the vaginal microbiota in this murine model with two primary objectives: A) to define how the murine vaginal microbiota changes throughout non-diabetic and diabetic pregnancy and B) to determine if there are signatures of vaginal communities that are protective or permissive against GBS *in utero* dissemination. Consistent with reports in nonpregnant mice^85,86^ the predominant taxa in pregestational C57BL/6J mice included *Staphylococcus*, *Lactobacillus*, *Enterococcus, Corynebacterium*, and *Streptococcus* spp. (**Fig. 7A,B, Fig. S5A,B**). We also detected *Enterobacteriaceae*, *Bacillus* and *Lactococcus* spp. (**Fig. 7A,B**). In pregnant controls, the vaginal microbiota increased in alpha diversity between the day prior to mating (d-1) and E14, whereas no change occurred in the GDM group (**Fig. 7C**). When we compared all premating timepoints (d-7 to d-1) to gestational or post-mating timepoints (d2-d14), increased alpha diversity was specific to pregnant control mice and absent in the non-pregnant cohort (**Fig. 7D-E, Fig. S5C-D**). This indicates a pregnancy-specific phenomena and confirms GDM-mediated disruption of pregnancy adaptation by the murine vaginal microbiota (**Fig. 7D-E**). Clustering by Bray-Curtis (BC) dissimilarity indices at pre-gestation, early gestation and mid-gestation revealed several key findings: A) that differences in vaginal communities were driven by 4 taxa (*Staphylococcus*, *Enterococcus*, *Lactobacillus,* and *Bacillus*), B) diabetic status did not impact overall vaginal community structure, and C) vaginal communities drifted away from *Staphylococcus succinus* dominance towards later points in gestation (E11, E14) in both non-diabetic and diabetic mice (**Fig. 7F-J, Fig. S5F-H**. *S. succinus* relative abundance significantly decreased by mid-gestation for all pregnant mice irrespective of diabetic status (**Fig. 7J**) but no changes occurred in non-pregnant mice (**Fig. S5K**). There were no GDM-mediated differences in intra-mouse pairwise BC dissimilarity between consecutive timepoints (**Fig. 7F**), but *Enterococcus* was a feature detected by an Analysis of Composition of Microbes (ANCOM) that was specific to GDM pregnancy (**Fig. 7K, Fig. S5I-J**). Additionally, ANCOM of samples from dams that had fetal GBS invasion vs. those that did not revealed that vaginal *Lactobacillus* and *Enterobacteriaceae* varied significantly between groups (**Fig. 7L**). Pregnant controls that had perinatal GBS invasion had significantly less *Lactobacillus* at mid-gestation (**Fig. 7M**). Increased *Enterobacteriaceae* during early gestation was associated with perinatal invasive disease in both pregnant controls and GDM groups (**Fig. S5L**). The GDM cohort had significantly greater relative abundance of *Enterococcus* on E2 suggesting an early enterococcal bloom (**Fig. 2N, Fig. S5E**), and by E17.5, a significantly greater proportion of GDM dams (44% vs. 12.5% of controls) experienced spontaneous uterine-fetal co-ascension by *Enterococcus* (**Fig. 7O**). Together, these analyses reveal that GDM uniquely alters vaginal microbiota dynamics in pregnancy and implicates specific taxa in influencing GBS perinatal disease.

**Figure 7:**
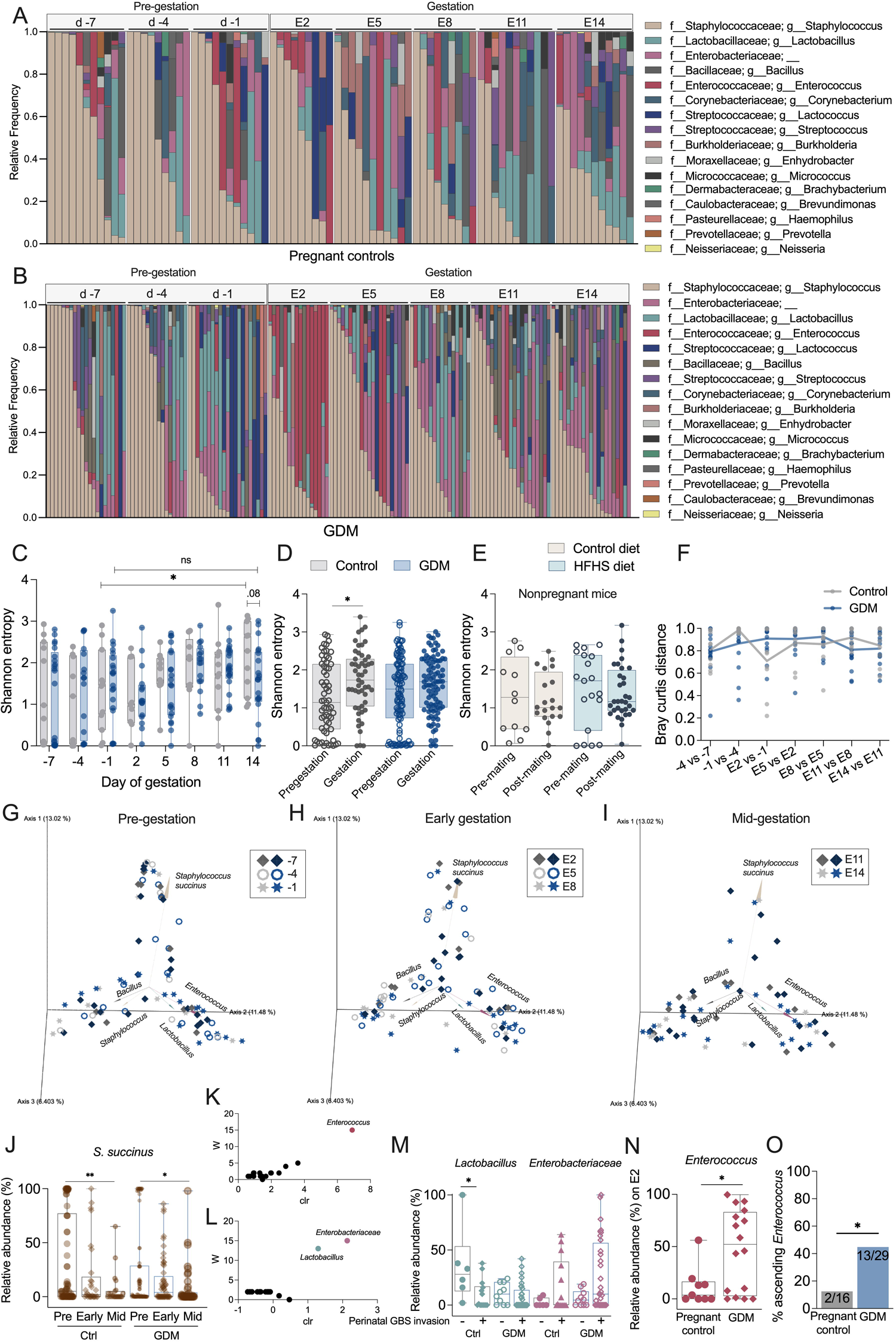
Pregnancy and gestational diabetes influence the vaginal microbiota composition and specific taxa correlate with GBS offspring dissemination. **A)** Pregnant control and **B)** GDM mice were swabbed every three days from one week before mating (d-7), until E14 before GBS challenge, and longitudinal vaginal microbiome composition was determined by 16Sv4 rRNA amplicon sequencing. **C)** Shannon entropy of vaginal communities at each timepoint before and throughout gestation. **D)** Composite Shannon entropy of all timepoints before gestation (d-7 through −1) and during gestation (E2 through E14) in pregnant controls vs. GDM mice. **E)** Composite Shannon entropy of all timepoints before mating vs. after mating in non-pregnant mice on control diet or high-fat high-sucrose (HFHS) diet. **F)** Bray Curtis distances between sequential timepoints per mouse for pregnant controls and GDM groups. Bray Curtis distance matrix principle coordinate analysis (PCoA) plots of vaginal communities at **G)** pre-gestation (d-7--1), **H)** early gestation (E2-E8), and **I)** mid-gestation (E11, E14) timepoints, with gray symbols representing pregnant control samples and blue symbols representing GDM samples. **J)** Relative abundance of *S. succinus* at pre-gestation, early gestation, and mid-gestation timepoints. **K)** Analysis of Composition of Microbes (ANCOM) in GDM mice across all timepoints. **L)** ANCOM in mice with GBS offspring dissemination (defined as GBS detected in fetal or pup livers) across all timepoints. **M)** Relative abundances of *Lactobacillus* and *Enterobacteriaceae* at mid-gestation timepoints, stratified by GBS dissemination. **N)** Relative abundance of *Enterococcus* on E2. **O)** Proportion of dams with *Enterococcus* ascension defined by detection in uterine, placental or fetal tissues. Points represent individual samples and lines indicate medians. Box and whisker plots extend from 25^th^ to 75^th^ percentiles and show all points (C-E, J, M-N). Data are from 4 independent experiments, *n* = 9-11 pregnant controls per timepoint, *n* = 16-21 GDM mice per timepoint, *n* = 3-8 non-pregnant mice on control diet per timepoint, *n* = 6-11 non-pregnant mice on HFHS diet per timepoint. For endogenous *Enterococcus* ascension, *n* = 16 control and 29 GDM dams aggregated from experiments shown in Figure 1. Data were analyzed by Welch’s t-test (C), Mann-Whitney t-test (D-E,M-N), Friedman’s tests with correction for multiple comparisons to assess differences between timepoints within each group (F), Kruskal-Wallis test with a post-hoc Dunn’s multiple comparisons test (J) and Fisher’s exact test (O). *p<0.05, ***p<0.001.

## DISCUSSION

Clinical data provide strong associations between diabetes in pregnancy and GBS vaginal carriage^33–36^ and neonatal disease^38^, but the molecular mechanisms driving these associations are undefined. Here, we describe the first animal model interrogating group B *Streptococcus* pathogenesis in gestational diabetes that mirrors the natural course of GBS vaginal colonization, *in utero* dissemination, and neonatal transmission. We observed that GDM mice had a significantly greater risk of GBS fetal infection and worse neonatal outcomes. We identified dysregulated maternal immunity, an altered maternal vaginal microbiota, and discordant GBS transcriptional adaptation including a novel GBS gene, *yfhO*, as mechanistic contributors to GBS disease in GDM hosts. Increased vaginal GBS burdens in GDM mice is consistent with reports of increased risk of GBS vaginal carriage in diabetic pregnant women (gestational and/or pregestational diabetes)^33–36^. Others found no association^87–89^ or associations solely when racial/ethnic backgrounds were considered^90^. These conflicting findings are possibly explained by GDM heterogeneity including varied severity of hyperglycemia and insulin dependency, immune dysfunction, GBS strain-dependent behavior, and other compounding biological and socioeconomic factors. Our model of genetically and phenotypically similar GDM hosts^59,60^ permitted mechanistic resolution of GBS maternal colonization and perinatal disease. Observed ranges in maternal and fetal burdens are likely related to the 2-3 day mating window used to maximize successful mating. Timing and dosage of GBS impact murine birth outcomes with earlier (E13, E14) and greater inocula associated with preterm birth^18,21^ compared to later (E17, E18) and lower inocula^75^. Despite the range in gestational age and two inoculations, we only observed 2 cases of preterm birth and both cases were in neutrophil-depleted GDM dams. Ultimately, the range in gestational age and tissue burdens provide a clinically relevant model reflective of GBS exposure patterns experienced by women.

Our data suggest that poor GDM pup outcomes (reduced survival and weight gain) were not driven by differences in postnatal GBS burdens, but perhaps were driven by an earlier disadvantage *in utero* shown by increased prenatal bacterial load. Our findings agree with a clinical report showing a 2-fold increased risk for early onset GBS sepsis or pneumonia for infants born to GDM mothers^38^. In humans, neonatal sex impacts GBS outcomes; male neonates with invasive GBS have a greater long-term risk of neurodevelopmental impairment and/or epilepsy^91,92^. We did not assess neonatal neurological development, but a sex dichotomy was detected at postnatal d7 with GDM males bearing higher invasive GBS burdens. Male pups have increased circulating IL-1β and placental IL-1β and CXCL1 in rat models of *in utero* GBS exposure^93^, but this was not replicated in our study (data not shown). We also assessed sex-specific differences *in utero.* Females in the control group displayed greater GBS invasion, with no sex differences in GDM fetuses. Intrauterine position effects might shape these findings; females positioned near males have an altered hormonal environment that bears long-term effects on anatomy and physiology^94^. Key aspects of our future work are delineating the contributions of fetal and neonatal immunity to perinatal outcomes, which we suspect are negatively impacted by GDM. Another critical future direction is exploring the neonatal microbiota in this model, as there is evidence of GDM-mediated dysbiosis in human neonates including increases colonization by *Streptococcaceae* and *Enterococcaceae*^53,95,96^, which may contribute to discrepant outcomes.

Previous studies have described GBS transcriptional adaptations to the murine non-gravid vaginal environment compared to *in vitro* culture conditions^31,32^. To our knowledge, this study provides the first comparison of GBS transcriptional responses between different tissues within the gravid reproductive tract with 22 DEGs identified between vaginal and uterine sites. There was minimal overlap between these genes and the prior studies with the exception of ribose metabolism (SAK_RS00825)^31^. Several DEGs overlapped with genes identified as critical for GBS colonization of the non-gravid murine reproductive tract using a GBS transposon mutant library including an acetyltransferase (SAK_RS07995), biotin synthase, a CHAP domain-containing protein (SAK_RS08980), and the peroxide-responsive transcriptional repressor *perR*^32^. We also detected significantly higher transcription of *mrvR* in vaginal tissues from pregnant controls; *mrvR* was recently reported to mediate nucleotide metabolism and provide GBS a growth advantage in amniotic fluid^29^. In high glucose media, GBS alters expression of genes involved in sugar transport, amino acid metabolism and transcription regulation^26^. In agreement, tissue-specific and group-specific transcriptional signatures of carbohydrate transport, metabolism, and transcription regulation in our study suggest metabolically unique environments within the reproductive tract. We suspect higher tissue glucose in GDM mice which would explain the upregulation of the GRP family sugar transporter (SAK_RS10655) and YfhO, a putative c-type glycosyltransferase, in control uterine tissues but not GDM. In this model, we found that *yfhO* was crucial for fetoplacental infection, with GDM conditions being permissive to this otherwise attenuated phenotype. The 2 putative stress response genes uniquely regulated in GDM tissues are yet to be characterized; however, the altered GBS-host interactions and increased pathogenicity in GDM is further supported by the upregulation of host stress, hypoxia and DNA damage pathways in GDM uteri.

Multiple clinical studies have identified aberrant systemic immune profiles in women with GDM including increased peripheral neutrophils, B cells, and CD8^+^ T cells^42,97,98^, a shift from CD56^bright^ (cytokine producing) NK to CD56^dim^ (cytotoxic) NK cells^44,49,50^. Reports of regulatory T cell differences are mixed^40,41,44,99^. Although we did not observe systemic differences across tissues in our model, our findings corroborate immune dysregulation with specificity to the maternal reproductive tract and in response to an opportunistic bacterial pathogen. A limitation of our study is that we did not immune profile mock-infected control and GDM mice and it remains possible that there are baseline immune differences in GDM mice; however, cytokine levels were quite similar in mock-infected controls suggesting minimal altered reproductive immune function in the absence of infection. Another limitation is that immune responses were evaluated 72h after GBS challenge, and thus more acute differences may have been missed. Characterizing other markers of aberrant inflammation such as immune cell sub-tissue localization and activation, and tissue damage, are key next steps.

Placentae from women with GDM have decreased regulatory T cells^100^, and increased NK cells^49^, neutrophil infiltration and activation^51^, and macrophage activation^101^. Our data suggest that placental susceptibility to GBS is in part driven by impaired immune recruitment during infection. Other potential contributing factors include altered immune cell activation, compromised placental integrity, or aberrant iron storage^102^, which have been reported in human GDM placentae^103,104^. We honed in on two cell types of interests arising from comparisons between GDM and control hosts: neutrophils, based on cytokine and host transcriptional differences, and natural killer cells, based on differing uterine proportions and placental counts in flow cytometry analyses.

With evidence of decreased uterine NK cells in GBS-challenged GDM mice, we hypothesized that NK depletion would worsen fetal infection. Surprisingly, NK depletion protected against GBS fetal invasion in GDM mice suggesting a deleterious role of NK cells during GBS infection in gestational diabetes. Although NK cells were systemically depleted, the observations that 1) infection was constrained to the reproductive tract in most mice (**Fig. 1C**), and 2) that the proportion of CD69^+^ NK cells was increased specifically in the uterus (**Fig. S4B**), together suggest observed effects were predominantly driven by uNKs, not peripheral NKs (pNKs). Murine and human uNKs are proangiogenic and less cytotoxic compared to pNKs^105,106^, constitutively express CD69 in mice^74^, and display receptors for MHC I and MHC ligand recognition, enabling cooperation with antigen presenting cells^105,107^. Studies on uNKs in local bacterial infections in pregnancy are scarce^107–109^; one study on decidual NKs (dNKs) showed that human dNKs selectively transfer granulysin to kill intracellular *Listeria monocytogenes* in placental cells^110^. NK depletion in bacteremic non-pregnant mice improves survival against some bacterial species^111,112^, worsens survival in some instances^113^, and does not appear to impact GBS systemic infection^108,111^. GBS itself may manipulate NK; GBS induces robust IFNy production in peripheral NKs *ex vivo* ^111^ and engages Siglec-7, a CD33-related inhibitory receptor, to suppress NK inflammasome activation^114^. How NK anti-GBS activity is affected by the maternal-fetal tolerogenic environment of pregnancy, or diabetic pregnancy, are unknown. Here, for the first time, we provide evidence that NKs are detrimental to control GBS ascending fetoplacental infection, and display decreased activation, in GDM hosts. We speculate that NK depletion in GDM mice contributed to a reduced inflammatory state and improved control of GBS fetoplacental invasion, and that the beneficial impacts of reduced NK cells were muted in the less inflammatory pregnant controls.

Neutrophils and macrophages are recruited to gestational tissues and engage various antibacterial strategies to limit ascending GBS infection^102,115,116^; however, GBS may evade immune responses by dampening activation^117^ or evasively hiding in macrophages^118^. Macrophage depletion decreases fetoplacental GBS burdens, but not maternal reproductive burdens, in a mouse model of ascending infection^118^. In this study, we are the first to report that, in addition to increasing vaginal GBS carriage, neutrophil depletion increases the frequency of GBS placental positivity without affecting fetal invasion, implying that other protective mechanisms constrain GBS in non-diabetic pregnancy. Neutrophil depletion had minimal impact on GBS pathogenesis in GDM mice, although it is possible that our study was underpowered to detect differences: we observed 40% incidence of preterm birth in neutrophil-depleted GDM dams and 0% in neutrophil-depleted controls but this difference was not significant. It is also possible that GDM mice are at the upper threshold or limit of GBS susceptibility, and thus it may not be biologically plausible in our model to further aggravate GBS infection.

In healthy human pregnancy, *Lactobacillus* spp. are enriched compared to non-pregnant or post-partum individuals and in general display more stability and lower alpha diversity^119–122^. Notably, the vaginal microbiome of pregnant mice is understudied despite the abundance of mouse models of obstetric disease. Three murine studies have characterized the maternal vaginal microbiota at discrete gestational timepoints and predominant vaginal taxa vary widely across studies^123–125^. This is the first longitudinal characterization of the vaginal microbiota in pregnant and diabetic pregnant mice, an important step in contextualizing the role of this bacterial community in reproductive health models. The dominant vaginal taxa in pregnant mice overlapped with non-pregnant mice from the same cohorts and from prior studies in non-pregnant C57BL/6J mice from Jackson labs^85,86,126^ suggesting that pregnancy does not alter the overall community composition, although we did detect reduction in *S. succinus* mid-gestation. In pregnant controls, but not GDM mice, we observed increased alpha diversity as gestation progressed, which has been reported in some women with *L. iners* and *L. crispatus* dominant communities^121^. The vaginal microbiota in women with GDM is more diverse and enriched for dysbiotic genera including *Bacteroides, Klebsiella, Enterococcus,* and *Enterobacter,* and consequently alters neonatal microbial inheritance in oropharyngeal and intestinal tissues^53,54,82,83^. Similarly, we observed increased *Enterococcus* in vaginal tissue in GDM mice and increased co-ascension of *Enterococcus* with GBS into the uterine and/or fetoplacental tissues in a subset of mice. Whether *Enterococcus* is a bystander or contributing to pathogenesis of these tissues is currently unknown. We also observed that control dams who experienced GBS invasion in offspring had reduced *Lactobacillus* during mid-gestation, while early gestation *Enterobacteriaceae* abundance increased in both GDM and controls that experienced perinatal invasive GBS. Together, these data demonstrate that this model recapitulates aspects of vaginal dysbiosis in gestational diabetic hosts and implicates specific taxa in influencing GBS pathogenesis in pregnancy. The functional roles of these taxa and whether differences in maternal microbiota composition instigate differences in neonatal microbial communities requires further investigation.

Our translational mouse model of GBS vertical transmission in gestational diabetic hosts recapitulates several clinical features and provides a foundation for further mechanistic and therapeutic exploration. Our findings highlight multifactorial drivers of GDM susceptibility to fetoplacental infection which include maternal immunity, pathogenic bacterial adaptations, and disruption of vaginal microbial changes in pregnancy. This preclinical model captures biological variability in GBS interactions with the gravid host that may aid in risk-stratification of women with GDM and provides a critical platform for testing alternative treatment options to improve perinatal outcomes in gestational diabetic pregnancy.

## MATERIALS AND METHODS

### Bacterial strains, growth conditions and inoculation preparation

GBS strains A909 (ATCC BAA-1138) and CNCTC 10/84 (ATCC 49447), or isogenic A909 Δ*cylE*^73^, or isogenic A909 Δ*yfhO,* were grown to stationary phase at 37°C in Todd-Hewitt broth (Hardy Diagnostics) for at least 16 h. Cultures were diluted in fresh Todd-Hewitt broth and incubated at 37°C until mid-logarithmic phase (defined as OD600 = 0.4-0.6). Bacterial cultures were centrifuged (3,220 *g*, 5 min), washed in sterile PBS, and resuspended at the desired concentration in sterile PBS.

### Construction of GBS *yfhO* mutant strain

To delete y*fhO* (SAK_RS10730, NCBI RefSeq accession NC_007432.1) in GBS A909, we used an allelic exchange approach as previously described^127^. Briefly, GBS A909 chromosomal DNA was used as a template for amplification of two 700bp DNA fragments using two primers pairs: YfhO-BamH-F(5’-CGT CTG GAT CCC TGC ACT TAT TGG ACA AAA TG-3’)/Kan-YfhO-R1(5’-CAG TAT TTA AAG ATA CCG GTA TAC GAA GCT TAT AGT G-3’) and Kan-YhfO-R2(5’-TGA TGA AAG CCA TCG CGT ACT AAA ACA ATT CAT CCA G-3’)/YfhO-XhoI-R(5’-GTG CGC TCG AGC TAC ATA AAT CAT AGG AAT AGA GCC-3’). The designed primers contained 16-20bp extensions complementary to the nonpolar kanamycin resistance cassette. The nonpolar kanamycin resistance cassette was PCR-amplified from pOSKAR (GenBank ID HM623914) using the primer pair YfhO-Kan-F1(5’-ATA AGC TTC GTA TAC CGG TAT CTT TAA ATA CTG TAG-3’)/YfhO-Kan-R2 which contained 16-20 bp extensions of complementary two DNA fragments of *yfhO*. The two fragments of *yfhO* and the fragment with the kanamycin resistance cassette were purified using the QIAquick PCR purification kit (Qiagen) and fused by Gibson Assembly (SGA-DNA) using primer pair YfhO-BamH-F/YfhO-XhoI-R. The assembled DNA fragment was digested with BamHI/XhoI and cloned into pHY304 digested with the respective enzymes. The plasmid was transformed into competent GBS A909 cells by electroporation and erythromycin resistant colonies were selected on THY agar plates at 30°C. Integration was performed by growth of transformants at 37°C with erythromycin selection. Excision of the integrated plasmid was performed by serial passages in THY media at 30°C and parallel screening for erythromycin-sensitive and kanamycin-resistant colonies. Double-crossover recombination was confirmed by PCR and Sanger sequencing using the following primer pair: YfhO-internal-F (5’-GGT TGG AAC AAA TAG TGT CC-3’) and YfhO-internal-R (5’-GAA TAA GCT GTT TGA ACC ATG-3’).

### Animals

Wild-type (WT) female and male C57BL/6J mice aged 6 weeks were purchased from Jackson Laboratories (strain code 000664). Male mice were used solely for mating and female mice were used for all *in vivo* experiments. Control and GDM groups were assigned randomly and mice were housed at 3 animals per cage. Mice ate and drank *ad libitum* and had a 12h light cycle per day. Timed matings were performed by introducing one male into a cage housing 3 females for a total of either two nights (pup outcomes experiments) or three nights (for *in utero* dissemination and immune profiling experiments). After each night, males were rotated between control and GDM cages to equalize exposure to productive male breeders. Soiled bedding from male cages was introduced into female cages three days before mating to synchronously induce estrus and promote successful mating. Embryonic day 0.5 (E0.5) of gestation was considered noon of the day after the first night of mating. All animal protocols and procedures were approved by the BCM Institutional Animal Care and Use Committee.

### *In vivo* model of GBS vaginal colonization in gestational diabetic mice

To establish gestational diabetes in mice, a diet-induced approach was used as previously described^59^. Briefly, mice are fed either a high-fat high-sucrose (HFHS) diet (D12451, Research Diets Inc.) or a control low-fat, no-sucrose diet (D12450K, Research Diets Inc.) one week before mating and maintained on the diet throughout pregnancy until the experimental endpoint. On days 14.5 and 15.5 of gestation, 1×10^7^ colony forming units (CFUs) of GBS in a 10μL PBS solution, was introduced into the vaginal tract with a gel loading pipette tip as described previously^128^ to model mid-gestational vaginal colonization by GBS. Due to the multi-day mating schedule as described above, there is a range in day(s) of GBS challenge of E12.5-E15.5 for mice mated for 3 nights and E13.5-E15.5 for mice mated for two nights. Accordingly, the experimental endpoint of E17.5 has a +/−2 or +/- 3 day window.

### GBS *in utero* dissemination and pup outcomes analyses

To assess dissemination of GBS from the vaginal inoculation site upwards to the uterine-fetal space before birth, mice were inoculated with GBS as described above and then sacrificed on day E17.5, two days before expected delivery. Maternal (vagina, cervix, uterus), fetal (liver), and maternal-fetal (placenta) tissues were harvested. Placentae and livers were collected from each conceptus per dam, thereby permitting comprehensive tracking of GBS *in utero* dissemination and placental-fetal invasion. Dissected tissues were placed into 2 mL screwcap tubes containing 500 μL PBS and 1.0mm zirconia/silica beads. The tubes were weighed and then kept on ice until homogenized in a Roche Magnalyser bead beater at 6000 rpm speed for 60 seconds. Ten-fold serial dilutions of tissue homogenates were plated on selective CHROMagar StrepB (DRG International, Inc.). Recovered GBS was identified as pink/mauve colonies and growth of blue colonies was considered endogenous *Enterococcus* spp. based on manufacturer protocols. Randomly selected blue colonies from vaginal, uterine and placental tissues were confirmed as *E. faecalis* by 16S rRNA gene sequencing. *In utero* litter features such as fetal intrauterine positions, fetal resorptions and total number of fetuses were also recorded. Dams with GBS detected in uterine and/or fetal samples were considered to have ascending infection. Offspring sex was determined by PCR with primers targeting *Rbm31x/y* F (5’-CAC CTT AAG AAC AAG CCA ATA CA-3’) and R (5’-GGC TTG TCC TGA AAA CAT TTG G-3’) as previously described^129^.

In addition to characterizing the susceptibility of gestational diabetic mice and their fetuses to ascending GBS infection before birth, we also assessed pup outcomes during the first week of life in a separate mouse cohort. Mice were inoculated as described above, individually housed after E15.5 inoculation and then monitored twice daily for pre-term labor, pup birth and survival until postnatal day 7. On postnatal days 3 and 7, pups were weighed. Upon death or postnatal day 7 sacrifice, pup intestines and livers were harvested and processed as described above to quantify GBS burden. All tissue homogenates were stored at −20°C until further use.

### Quantifying cytokine levels in vaginal, uterine and placental tissues

Vaginal, uterine, and placental tissue homogenates cytokines (IL-1α, IL-1β, IL-2, IL-3, IL-4, IL-5, IL-6, IL-9, IL-10, IL-12 (p40), IL-12 (p70), IL-13, IL-17A, Eotaxin, G-CSF, GM-CSF, IFN-γ, KC, MCP-1, MIP-1α, MIP-1β, RANTES, and TNF-α) were quantified via a 23-plex assay (Bio-Rad, Cat. No M60009RDP). Tissue samples were thawed on ice and then spun at 10,000*g* for 10 minutes. Vaginal and uterine tissue supernatant were diluted 1:10 in Bio-Rad sample diluent, and placental tissues were diluted 1:2. Samples were then processed per the manufacturer’s protocol. Data was acquired with Luminex xPONENT for Magpix, version 4.2 build 1324 on a Magpix instrument, and data was analyzed with Milliplex Analyst, version 5.1.0.0.

### Immune cell profiling of vaginal, uterine and placental tissues

Vaginal and placental tissues were transected with one-third processed for burden quantification and the remaining two-thirds processed for flow cytometric quantification of immune cell populations. For uterine tissues, each horn was transected in half, and the halves from each horn were pooled and processed for either burden quantification as described above, or for immune cell quantification. Tissues harvested for flow were placed in 450 μL (placenta) or 900 μL (vagina, uterus) of RPMI, and mechanically disrupted with scissors until ∼90% of sample was fine enough to pass through a p1000 tip. Collagenase (0.2mg/mL) and DNase (50 U/mL) were added to each vaginal and uterine sample, and Collagenase (0.2mg/mL) and DNase (50 U/mL) were added to placental samples. After vortexing, the samples were incubated at 37 °C and 250rpm shaking for 30 minutes. 350 μL of supernatant was then filtered (40 μm) into an Eppendorf containing 800 μL RPMI + 10% FBS, and the resulting filtered cells were kept on ice. The remaining tissue fragments underwent a second digestion after supplementing with 350 μL total of collagenase, DNase and RPMI at concentrations specified above per tissue type. After a second incubation at 37°C and 250rpm for 30 minutes, 350 μL of supernatant was filtered into the collection tubes containing single cells from previous filtration step. Samples were then spun at 500×*g* for 10 minutes, resuspended in 500 μL of Red blood cell lysis buffer (Lucigen, SS000400-D2) and incubated at room temperature for 5 minutes. 700 μL of PBS was then added, samples were spun at 500×*g* for 10 minutes, and then resuspended in 50 μL of PBS. Cells were then stained with 50 μL of Zombie Aqua that was reconstituted and diluted (1:1000 dilution) per manufacturer instructions. Cells were then incubated for 15 minutes at room temperature in the dark. Subsequently, cells were washed with 150 μL PBS, spun for 10 minutes and then resuspended in 50 μL of a 1:200 dilution of Fc block (CD16/CD32, clone 2.4G2, BD Biosciences, cat. 553141, 0.5mg/mL) in FACS buffer (PBS, 1mM EDTA, 1% FBS, 0.1% sodium azide), followed by a 15 minute incubation at 4°C in the dark. Cells were then incubated with an 18 antibody cocktail for 30 min, in the dark, at 4°C. The antibody cocktail comprised: anti-CD3 (BV480, clone 145-2C11, cat. 746368, BD Biosciences), anti-CD45 (BV605, clone 30-F11, cat. 563053, BD Biosciences), anti-CD11b (APC-Cy7, clone M1/70, cat. 561039), anti-CD11c (BV785, clone N418, cat. 117335, BioLegend), anti-Ly6G (AF700, clone 1A8, cat. 561236, BD Biosciences), anti-NK1.1 (BUV737, clone PK136, cat. 741715, BD Biosciences), anti-CD19 (BB515, clone 1D3, cat. 564509, BD Biosciences), anti-CD8a (PE-Cy7, clone QA17A07, cat. 155018, BioLegend), anti-CD4 (BUV563, clone GK1.5, cat. 612923, BD Biosciences), anti-CD23 (BUV395, clone B3B4, cat. 740216, BD Biosciences), anti-CD44 (BUV805, clone IM7, cat. 741921, BD Biosciences), anti-CD64 (BV421, clone X54-5/7.1, cat. 139309, BioLegend), anti-MHCII (BV650, clone M5/114.15.2, cat. 563415, BD Biosciences), anti-CD24 (BV711, clone M1/69, cat. 563450, BD Biosciences), anti-CD62L (PE, clone M1/69, cat. 161204, BioLegend), anti-CD69 (PE-CF594, clone H1.2F3, cat. 562455, BD Biosciences), anti-Ly-6C (PerCP-Cy5.5, clone HK1.4, cat. 128012, BioLegend), anti-CD25 (APC, clone PC61, cat. 557192, BD Biosciences). 0.25 μL of antibody was added to each sample, for all antibodies except for anti-CD23, anti-CD24, and anti-Ly-6C for which 0.125 μL was added, and anti-CD3 for which 0.5 μL was added. Antibodies were added to staining buffer (10 μL brilliant stain buffer, and staining buffer (cat., vendor) to bring volume up to 50 μL), for a total volume of 50 μL per sample, which was then added to all samples. After a 30 minute incubation, samples were then washed with 100 μL of PBS, spun down (500×*g*) for 10 minutes and then resuspended in about 300 μL FACS buffer. Data were acquired using a BD FACSymphony A5 and post-acquisition analyses were done using FlowJo software version 10.8. Gating strategy is shown in Fig 4. Immune cell subsets were delineated from the CD45+ Zombie aqua-population and defined based on the following staining profiles: basophils (CD19^−^, Ly6G^−^, MHCII^−^, CD11b^+^, CD62L^−^, Ly6c^+/−^ CD25^+^), dendritic cells (CD11b^+^, CD11C^+/−^, MHCII^+^, CD24^+^), eosinophils (CD19^−^, Ly6G^−^, MHCII^−^, CD11b^−^, CD62L^−^, Ly6c^lo^ CD25^−^), macrophages (CD11b^+^, CD11C^+/−^. MHCII^+^, CD24^+/-,^ CD64^+^), mast cells (CD19^−^, Ly6G^−^, MHCII^−^, CD11b^−^, CD62L^+/−^, Cd11c^lo^, SSCA^lo^), monocytes (CD19^−^, Ly6G^−^, MHCII^−^, CD11b^+^, CD62L^+^), neutrophils (CD19^−^, Ly6G^+^), natural killer cells (CD19^−^, Ly6G^−^, NK1.1^+^), MHCII^+^ other cells (CD19^−^, Ly6G^−^, CD11b^+^, CD11c^+^, MHCII^+^, CD24L^−^ CD64L^−^), MHCII^−^ other cells (CD19^−^, Ly6G^−^, CD11b^−^, CD11c^+^, MHCII^−^, CD62L^+/−^), B cells (CD19^+^, Ly6G^−^), CD4^+^ T cells (CD19^−^, Ly6G^−^, CD11b^−^, CD11c^−^, CD4^+^), regulatory T cells (Tregs) CD4_+_ T cells (CD19^−^, Ly6G^−^, CD11b^−^, CD11c^−^, CD4^+^, CD25^+^), active CD4^+^ T cells (Tregs) CD4_+_ T cells (CD19^−^, Ly6G^−^, CD11b^−^, CD11c^−^, CD4^+^, CD69^+^), naïve CD4^+^ T cells (CD19^−^, Ly6G^−^, CD11b^−^, CD11c^−^, CD4^+^, CD62L^+^), memory CD4^+^ T cells (CD19^−^, Ly6G^−^, CD11b^−^, CD11c^−^, CD4^+^, CD44^+^), CD8^+^ T cells (CD19^−^, Ly6G^−^, CD11b^−^, CD11c^−^, CD8^+^), naïve CD8^+^ T cells (CD19^−^, Ly6G^−^, CD11b^−^, CD11c^−^, CD8^+^, CD62L^+^), memory CD4^+^ T cells (CD19^−^, Ly6G^−^, CD11b^−^, CD11c^−^, CD8^+^, CD44^+^).

### Immune cell depletion experiments

For NK cell depletion experiments, mice received intraperitoneal injections of 250 μg of anti-NK1.1 antibody (Bioxcell, clone PK136), or 250 μg of mouse IgG2a isotype control (Bioxcell, clone C1.18.4) on E13.5 and E15.5. For neutrophil depletion experiments, mice received intraperitoneal injections of 200 μg of anti-Ly6G antibody (Bioxcell, clone PK136), or 200 μg of rat IgG2a isotype control (Bioxcell, clone C1.18.4) on E13.5 and E15.5. Mice were challenged with vaginal inoculations of GBS on E14.5 and E15.5 and sacrificed on E17.5 to assess *in utero* bacterial dissemination as described above. Uterine tissues were harvested and processed for flow cytometry as described above, with a few alterations in work flow: single cells were fixed in 2% paraformaldehyde in RPMI for 10 minutes, washed with 500 μL RPMI, and then stained with live/dead, CD45, CD11b, NK1.1. or Ly6G antibodies overnight at 4°C. Data were acquired and analyzed as described above to confirm targeted depletion.

### RNA-sequencing tissue processing and analysis

Vaginal and uterine tissues (E17.5) from GBS-challenged GDM and pregnant control mice were dissected and placed in tubes containing 500 μL RNA Protect and 1.0mm zirconia/silica beads. Vaginal tissues were transected with half processed for RNA-sequencing and half processed for GBS quantification. Left uterine horns were processed for GBS quantification and right uterine horns were processed for RNA sequencing. Tissues were kept on ice until homogenized in a Roche Magnalyser bead beater at 6000 rpm speed for 60 seconds, spun at 13,000 rpm for 10 min. The resulting supernatant was discarded and pellets were resuspended in 100 μL TE and stored at −80°C, until shipment to SeqCenter where RNA extraction, rRNA depletion and dual-RNA sequencing on an Illumina Stranded RNA-seq platform were performed. For bacterial sequences, quality control and adapter trimming was performed with bcl2fastq(bcl2fastq, #164), read mapping was performed with HISAT2^130^, and read quantification was performed using Subread’s featureCounts^131^, with alignment to the A909 reference genome. For murine sequences, quality control and adapter trimming were performed with bcl-convert^132^. Read mapping was performed via STAR^133^ to the mm10 version of the mouse genome and feature quantification was performed using RSEM^134^. Raw counts normalization and differential expression analyses were performed using R^135^ package DESeq2 (v 1.40.1)^136^. RStudio^137^ (2022.12.0+353) was used to generate all heatmaps, PCA and volcano plots and pathways enrichment plots. The enhanced volcano^138^ R package was used as well as ashr^139^ LFC shrinkage for data visualization. The fGSEA R package (v1.20.0, RRID:SCR_020938) was used for gene set enrichment analyses with 10,000 permutations, a minimum gene set of 15, a maximum gene set of 500, and the Reactome gene set collections from the Molecular Signatures Database^140,141^.

### YfhO structural predictions and protein-protein interactions

AlphaFold2^69,70^ was used to generate predictions from amino acid sequences for YfhO (*Streptococcus agalactiae* gene locus GL192_08955), transmembrane domains predicted by UniProt (A0A8I2JTU5)((Consortium, 2023 #162)) and image generated using ChimeraX 1.6.1^142^. Lastly, the YfhO protein sequence was uploaded into STRING^71^ (v 11.5), using strain 2603V/R as a reference, to predict proteins that are functionally associated with YfhO in GBS. Predicted associations required a medium-high interaction score of 0.65 or greater.

#### Longitudinal tracking of the maternal vaginal microbiota via 16S rRNA sequencing

To characterize the murine vaginal microbiota throughout healthy and gestational diabetic pregnancy, we serially sampled the vaginal lumen every three days beginning seven days before mating until day of GBS challenge (E14.5). Vaginal swabs were collected as described previously^107^. For each swab collection, we included an environmental control comprised of a swab exposed to the laminar hood air circulation for several seconds with subsequent submersion in sterile PBS. These PBS blanks were used as controls for environmental and sequencing contamination. Swab samples were then vortexed to dislodge biomaterial from the swab tips and the swabs were removed and discarded. The remaining PBS sample was then stored at −20°C until further processing. Bacterial DNA was extracted from thawed PBS swab samples with the Quick-DNA Fungal/Bacterial Microprep kit following the manufacturer’s protocol (Zymo Research) with final elution in 20 μL of water. The v4 region of the bacterial 16S rRNA gene was amplified by PCR using primer 515F and 806R and sequenced on an Illumina MiSeq v2 using the 2×250bp paired-end protocol by the BCM Alkek Center for Metagenomic and Microbiome Research (CMMR). Raw data files were converted into FASTQs and demultiplexed using the Illumina ‘bcl2fastq’ software and single-index barcodes. Demultiplexed read pairs underwent initial quality filtering using bbduk.sh (BBMap version 38.82, 5) removing Illumina adapters, PhiX reads and reads with a Phred quality score below 15 and length below 100 bp after trimming. Quality controlled reads were merged using bbmerge.sh (BBMap version 38.82, 5), and further filtered using vsearch (6) with a maximum expected error of 0.05, maximum length of 254 bp and minimum length of 252 bp. All the reads were then combined into a single fasta file by CMMR for further processing. We then joined and trimmed raw sequences to 150 bp, and denoised using Deblur through the DADA2 plugin on QIIME2 v2021.11 with taxonomic assignments determined using the naïve bayes sklearn classifier trained on the GreenGenes OTUs database (13_8, 99% sequence similarity). Dataset decontamination included filtering out OTUs that appeared in fewer than 5% of samples, removal of known DNA extraction contaminants^143–145^, and processing samples through the Decontam R package^146^. Qiime2 was then used for diversity and compositional analyses of filtered, unrarefied, reads.

### Statistical analyses

Statistical analyses were performed using GraphPad Prism v9.5.1 as described in figure legends. Briefly, normality was assessed by the D’Agostino-Pearson normality test. Analysis of non-parametric data included two-tailed Mann-Whitney T tests for two groups, or Kruskal-Wallis test with correction for multiple comparisons for three or more groups. When appropriate, post-hoc Dunn’s multiple comparisons test was utilized, or the two-stage linear step-up procedure of Benjamini, Krieger and Yekutieli to correct for multiple comparisons by controlling the false discovery rate (<0.05). Two-sided Fisher’s exact tests were used for contingency analyses. The log-rank (Mantel–Cox) test was used to analyze survival curves, with Holm-Šídák correction for multiple comparisons when appropriate. For multiple paired samples (longitudinal bray Curtis distance comparisons), analyses were conducted with Friedman’s tests and Dunn’s post hoc correction for multiple comparisons. Differentially expressed genes were determined with a generalized linear model and genes with a Log_2_ Fold Change >1 and FDR adjusted p-value < 0.05 were considered significantly different. For all experiments, a p-value of < 0.05 was considered significant.

## ACKNOWLEDGEMENTS and FUNDING

This project was supported by the Cytometry and Cell Sorting Core at Baylor College of Medicine with funding from the CPRIT Core Facility Support Award (CPRIT-RP180672), the NIH (CA125123 and RR024574) and the assistance of Joel M. Sederstrom, Padmini Narayanan and Claude Chew. The Center for Metagenomics and Microbiology Research for 16S sequencing (CMMR) and the Texas Medical Center Digestive Diseases Center (DDC), with funding from the NIH (DK056338), were also important resources for this study. We would also like to acknowledge Susan F. Venable from the DDC for assistance with the multiplex ELISAs, Simone R. Hernandez for acquisition of murine tissue samples, and Anaid Reyes from the CMMR for assistance with 16S sequencing. We are also thankful to Misu A. Sanson-Iglesias and Luis A. Vega for helpful suggestions and resources for dual-RNA sequencing experiments.

MEM and JJZ were supported by an NIH T32 award (T32GM136554) and VME, MEM, and SO were supported by NIH F31 awards (AI167547, AI167538, and HD111236) respectively. VME was also supported by a scholarship from Baylor Research Advocates for Student Scientists (BRASS) and a Grant for Emerging Researchers/Clinicians Mentorship (GERM) Program from the Infectious Diseases Society of America (IDSA). Studies were supported by the Burroughs Wellcome Fund Next Gen Pregnancy Initiative (NGP10103), NIH R01 (DK128053), and U19 (AI157981) to KAP.

## AUTHOR CONTRIBUTIONS

KAP and VME conceived and designed experiments. VME, MEM, JJZ, SO, ZH, CS, CMR, MGM and MBB performed experiments. NK, KAP, and ARF provided reagents. KAP and VME analyzed data and interpreted results. KAP, KAP, and ARF secured funding. VME and KAP drafted the manuscript. All authors reviewed and edited the manuscript.

