## Supplemental Material for "Gestational diabetes augments group B *Streptococcu*s perinatal infection through disruptions in maternal immunity and the vaginal microbiota"

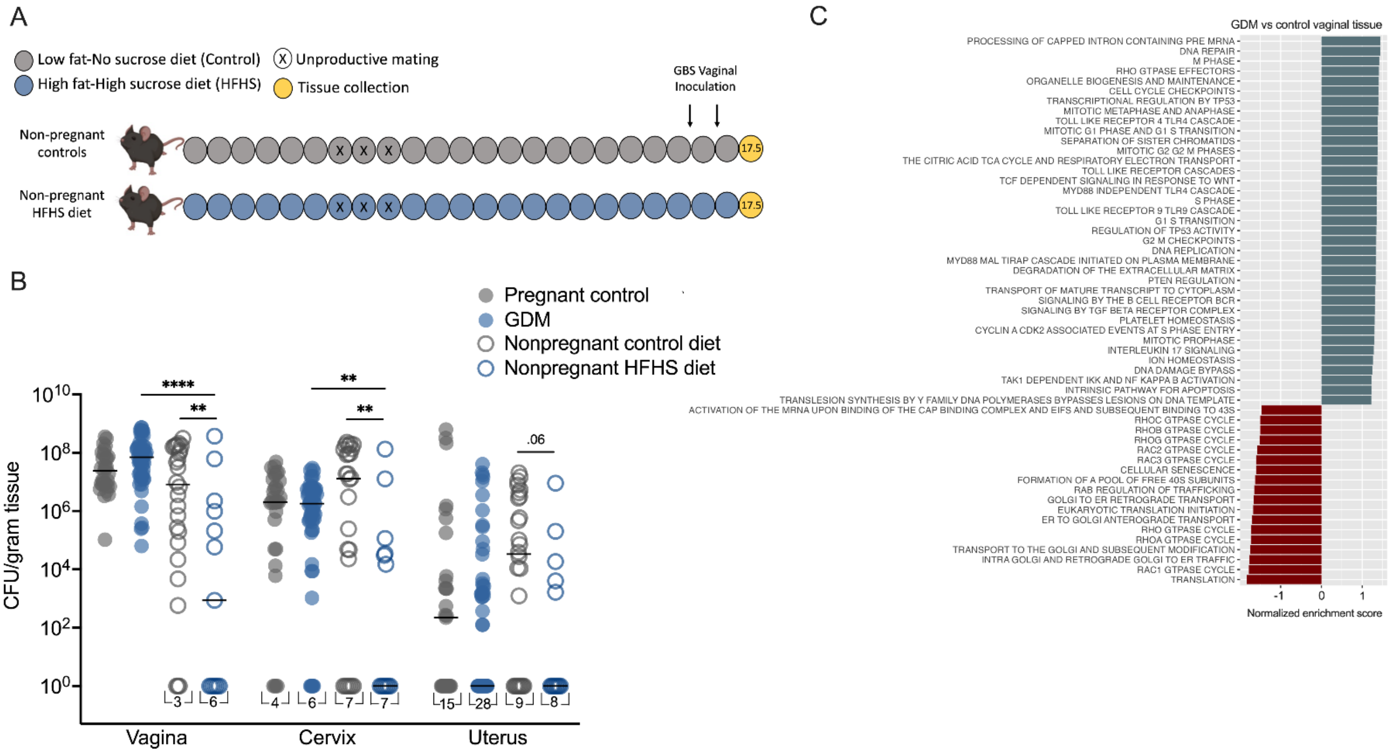


**Figure S1: Pregnancy and diet differentially impact GBS reproductive tract burdens, and gestational diabetes alters host transcriptional profiles in the vagina. A)** Parallel experimental timeline for non-pregnant mice maintained on either control or high-fat high-sucrose (HFHS) diet followed by vaginal colonization with GBS strain A909 on d14.5 and 15.5 and sacrifice on d17.5 to assess GBS dissemination. **B)** GBS burden throughout the reproductive tract where points represent individual mouse samples and lines indicate median CFU per gram of tissue. *n* = 31 pregnant controls, *n* = 54 GDM, *n* = 26 non-pregnant mice on control diet, *n* = 13 non-pregnant mice on GDM diet. Data were analyzed by multiple Mann-Whitney t-tests, **p<0.01, ****p<0.001. **C)** Gene set enrichment analysis of vaginal tissue in gestational diabetic mice vs. pregnant controls (related to **Fig. 3**).

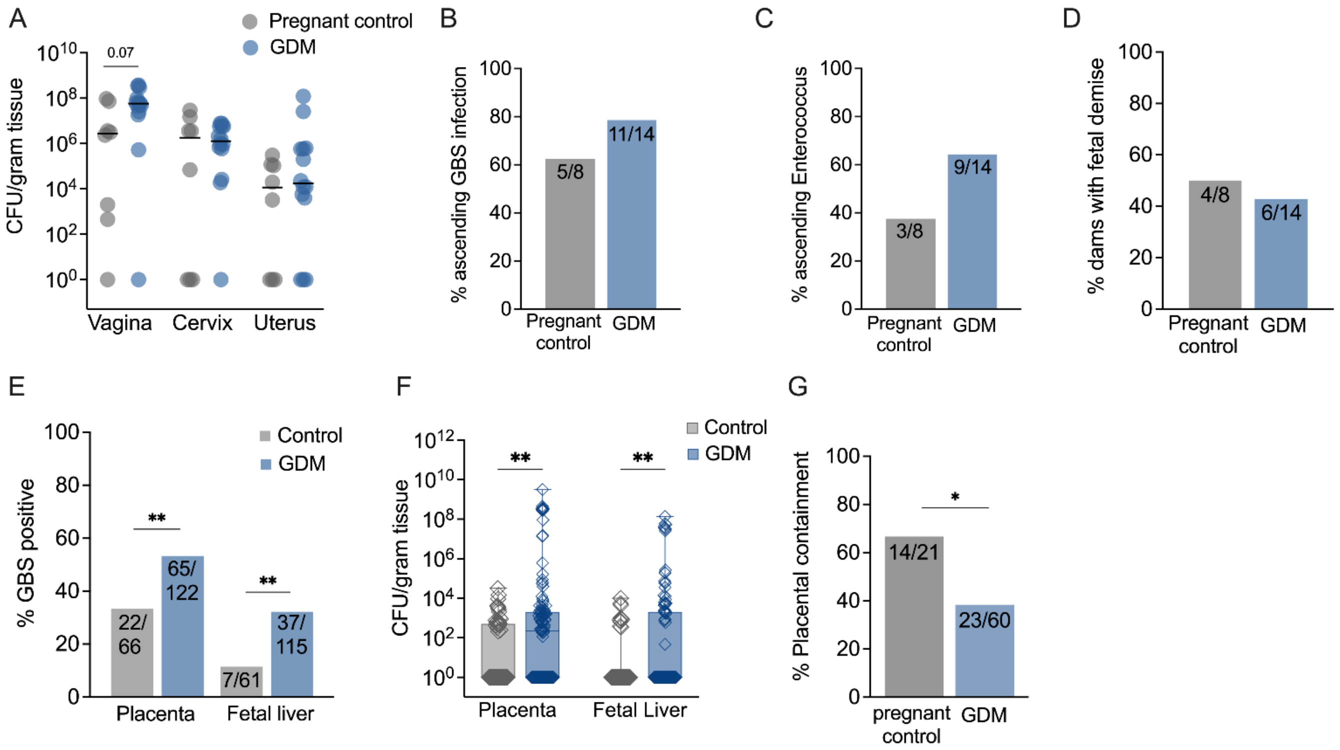


**Figure S2: Enhanced susceptibility of gestational diabetic mice to *in utero* group B Streptococcal fetal invasion is replicated by a GBS serotype V strain.** GDM was induced via a high-fat high-sucrose (HFHS) diet followed by mid-gestational GBS vaginal colonization with GBS CNCTC 10/84 and tissue collection on E17.5. **A)** GBS burden in maternal reproductive tract tissues. Proportion of dams with **B)** ascending GBS infection, **C**) ascending endogenous *Enterococcus*, or **D)** fetal demise (e.g. reabsorption). **E)** Percentage of placentae and fetal livers that were GBS positive, and **F)** corresponding GBS burdens. **G)** Percentage of placental-fetal units that had GBS detected in the placenta with no detection in the corresponding fetal liver. All data represent 3 independent replicates. Points represent individual samples and lines indicate medians (A,F). Box and whisker plots extend from 25$^{th}$ to 75$^{th}$ percentiles and show all points (F). Experimental numbers are $n$ = 8 control and 14 GDM dams, and experimental numbers for placenta-fetal pairs in each group are given as denominators in E and G. Data was analyzed by Mann-Whitney t-test (A,F) and Fisher's exact test (B-D, E, G). **p<0.01.

**Table S1: Differential gene expression of GBS in murine uterine vs. vaginal tissue.** Related to Figure 2.

| Current gene locus | Former locus tag | Gene name | Expected product | Pregnant control $log_2$ Fold-Change (*p*-value)[a] | GDM $log_2$ Fold-Change (*p*-value)[b] |
|---|---|---|---|---|---|
| SAK_RS07995 | SAK_1585 | | acetyltransferase | 3.61 (9.07E-34) | 3.95 (1.62E-43) |
| SAK_RS10655 | SAK_2115 | | GRP family sugar transporter | 3.57 (0.025) | NA |
| SAK_RS00885 | SAK_0179 | | L-lactate dehydrogenase | 2.83 (4.76E-12) | 3.17(2.90E-17) |
| SAK_RS10730 | SAK_2130 | yfhO | YfhO family protein | 2.39(0.048) | NA |
| SAK_RS09065 | SAK_1801 | rsmA | ribosomal RNA small subunit methyltransferase A | 2.37 (7.45E-07) | 2.55 (5.35E-09) |
| SAK_RS00825 | SAK_0167 | rbsC | ribose ABC transporter permease | 2.16 (1.77E-17) | 2.28 (1.15E-21) |
| SAK_RS02820 | SAK_0566 | bioB | biotin synthase | 1.87 (1.20E-05) | 1.88 (9.87E-06) |
| SAK_RS02870 | SAK_0576 | | DUF3165 family protein | 1.75 (0.044) | NA |
| SAK_RS08980 | SAK_1784 | | CHAP domain-containing protein | 1.38 (0.00005) | 1.28 (0.018) |
| SAK_RS08695 | SAK_1729 | | hypothetical protein | -1.31 (0.0004) | -1.23 (0.042) |
| SAK_RS01760 | SAK_0355 | proB | glutamate 5-kinase | -1.34 (0.0002) | -1.318 (0.0010) |
| SAK_RS10120 | SAK_2012 | mrvR | GntR family transcriptional regulator | -1.34 (2.86E-07) | NA |
| SAK_RS08640 | SAK_1718 | | DUF1129 family protein | -1.37 (0.028) | NA |
| SAK_RS08460 | SAK_1680 | gatA | glutamyl-tRNA(Gln) amidotransferase subunit A | -1.44 (2.86E-07) | NA |
| SAK_RS09300 | SAK_1847 | | N-acetyltransferase | -1.45 (1.20E-06) | NA |
| SAK_RS02315 | SAK_0465 | perR | peroxide-responsive transcriptional repressor | -1.47 (0.005) | -1.56 (0.00022) |
| SAK_RS01595 | SAK_0323 | | hypothetical protein | -1.71 (0.018) | -2.05 (4.50E-06) |
| SAK_RS07010 | SAK_1393 | | ABC transporter ATP-binding protein | -1.80 (3.74E-18) | -1.62 (2.09E-10) |
| SAK_RS06220 | SAK_1240 | | DUF2969 domain-containing protein | -2.51 (1.30E-31) | -2.46 (9.07E-30) |
| SAK_RS06520 | SAK_1300 | | DUF3042 family protein | -2.96 (4.75E-87) | -2.77 (4.36E-71) |
| SAK_RS04330 | SAK_0866 | | DUF3270 domain-containing protein | NA | 1.33 (0.049) |
| SAK_RS10715 | SAK_2127 | | YoaK family protein | NA | -1.50 (0.005) |

a. Fold-change between GBS isolated from uterine tissue vs. GBS isolated from vaginal tissue in pregnant control mice
b. Fold-change between GBS isolated from uterine tissue vs. GBS isolated from vaginal tissue in GDM mice

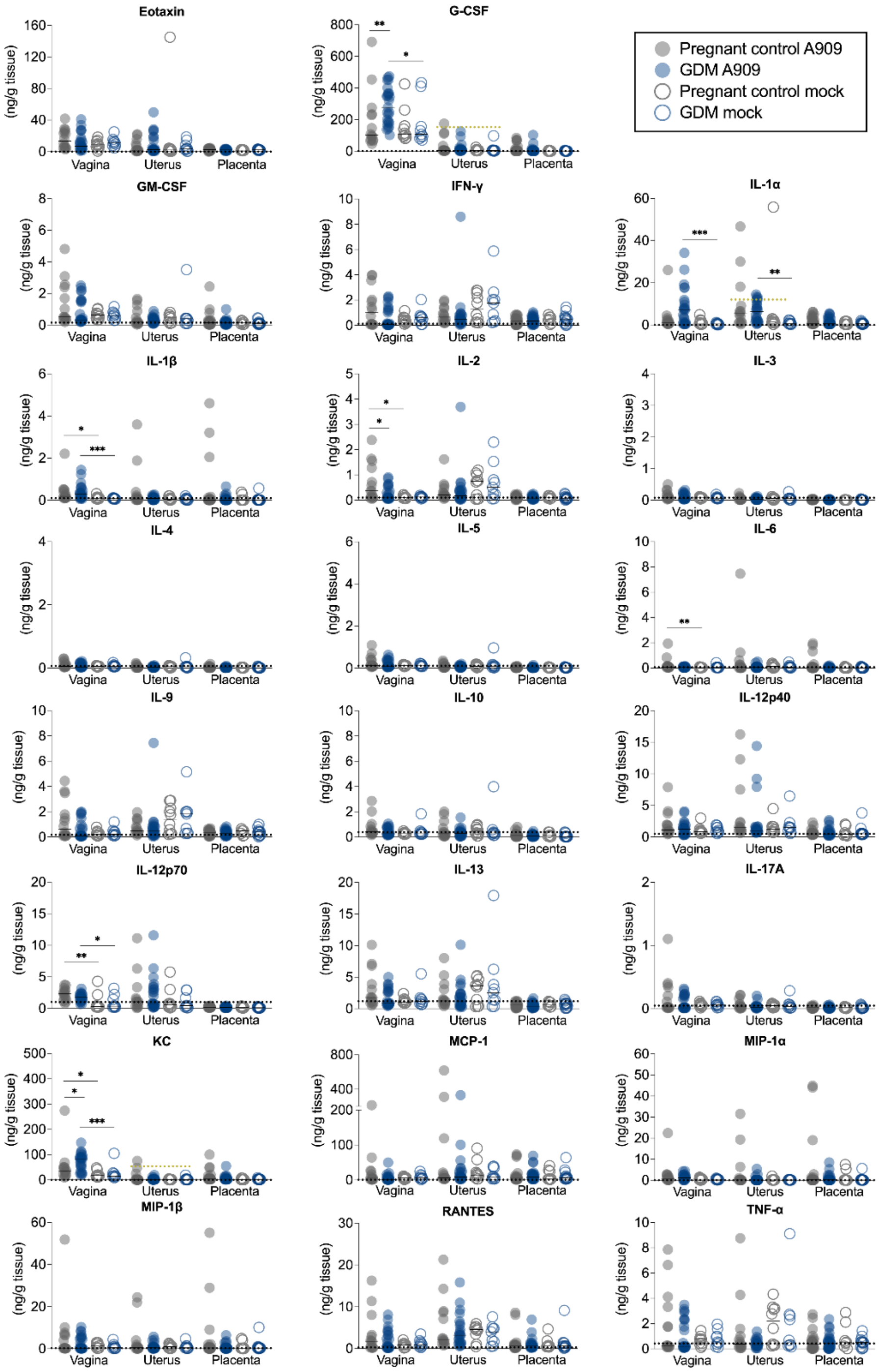


**Figure S3: Cytokine values from vaginal, uterine, and placental tissues.** Quantification of 23 cytokines in tissues on E17.5 from pregnant controls and GDM mice that were inoculated with A909 or mock-infected, all of which are presented as a heatmap in Fig. 3E. Each point represents an independent mouse sample and data were analyzed by Kruskal-Wallis test followed by a two-stage linear step-up procedure of Benjamini, Krieger and Yekutieli to correct for multiple comparisons by controlling the false discovery rate (<0.05). Black dashed lines demarcate lower limit of detection, and yellow dashed lines indicate upper limit of detection (ULD) when sample(s) surpassed. *p<0.05, **p<0.01, ***p<0.001.

**Table S2: DGEs of GDM vs. pregnant control reproductive tissues.** Related to Figure 3.

| Gene | Expected function(s) | $\log_2$ Fold-Change (*p*-value) |
|---|---|---|
| **GDM vaginal vs. pregnant control vaginal tissue** | | |
| Chil4 | Chitin binding activity and Chitin catabolic processes; Kinase binding activity; Positive regulator of chemokine production. | -6.95 (7.20E-04) |
| H2-Q6 | Antigen processing and presentation of endogenous peptide antigens via MHC class I and Ib. Positive regulator of T cell mediated cytotoxicity. | -21.92 (7.20E-04) |
| Eif3j1 | Member of eukaryotic translation initiation factor 3 complex. | -24.41 (2.63E-04) |
| Sbpl | Predicted to have activity in extracellular space. Orthologous to human zymogen granule protein 16B which is predicted to mediate carbohydrate binding. | -37.55 (7.29E-14) |
| **GDM uterine vs. pregnant control uterine tissue** | | |
| Scgb2b27 | Binds androgens; also known as Androgen binding protein beta (Abpb). Some secretoglobins have immunomodulatory functions. | 14.76 (9.90E-05) |
| Cxcl2 | Immunoregulatory and inflammatory processes. Chemotactic for polymorphonuclear cells and hematopoietic stem cells. | -7.46 (1.67E-02) |
| Odam | Involved in wound response mechanisms. | -15.54 (4.00E-04) |
| Rad21l | Chromatin binding activity; double-strand break repair via homologous recombination; homologous chromosome segregation. | -21.0 (5.10E-03) |
| Nlrp14 | Predicted to enable ATP binding activity and involved in cell differentiation. May play a role in inflammation as other members of the NALP protein family are immunomodulatory. | -21.47 (3.33E-03) |
| Prol1 | Implicated in stability of murine oral microbiome. Undefined function in uterus. | -21.62 (2.82E-06) |
| Zp2 | Encodes glycoproteins important for survival of growing oocytes, fertilization, and the passage of early embryos through the oviduct. | -21.88 (2.35E-03) |
| Padi6 | Protein-arginine deiminase activity Human ortholog(s) implicated in infertility. | -22.27 (2.87E-17) |
| Cst8 | Predicted to negatively regulate peptidase activity Expression specific to the reproductive tract, suggesting a role in reproduction. | -24.65 (9.90E-05) |

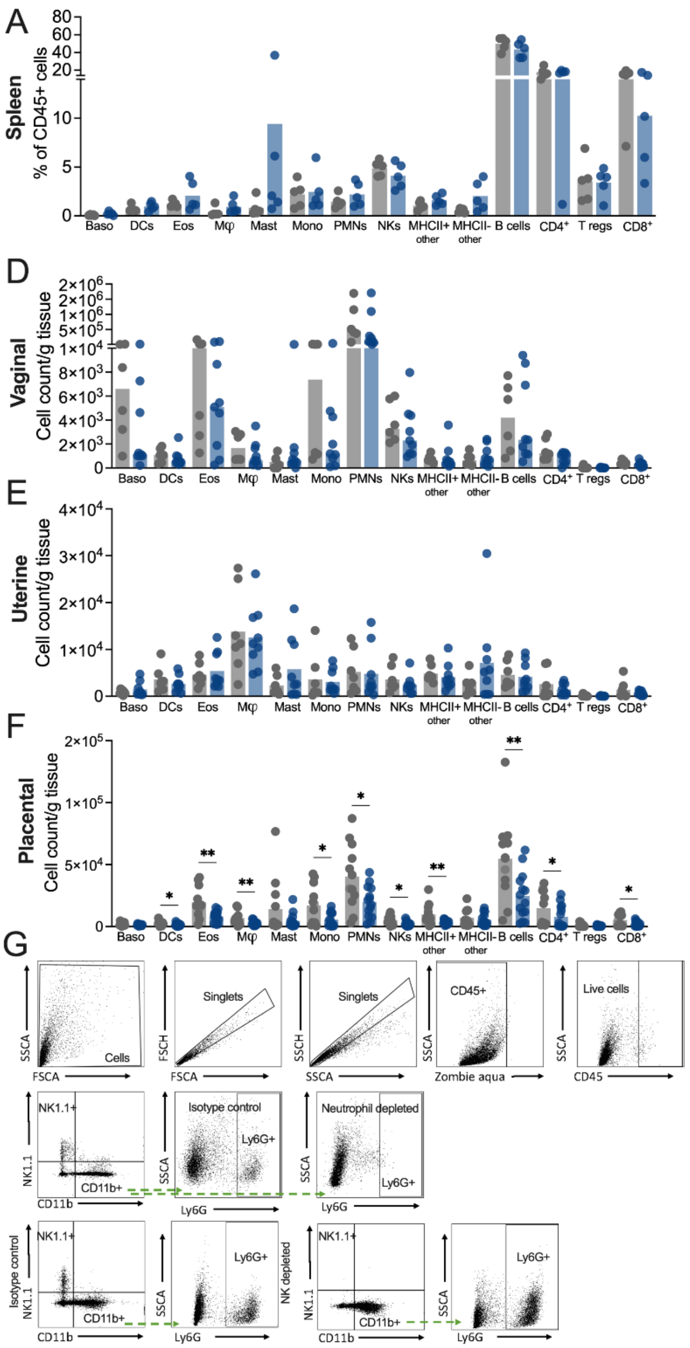


**Figure S4: Additional data for *in vivo* immune cell experiments. A)** Immune cell frequencies in splenic tissues from GBS-infected dams. **B)** Total cell counts and **C)** total live CD45+ cell counts per tissue normalized to tissue weight. Immune cell counts in **D)** vaginal, **E)** uterine, and **F)** placental tissues from GBS-infected dams. **G)** Gating strategy for assessing depletion of NK cells or neutrophils with α-NK1.1 and α-Ly6G antibodies respectively. Data (A-F) are from 3 independent experiments with each point representing an individual mouse sample (*n* =7 pregnant controls and 9 GDM, with 1-2 placentae per dam for a total of *n* = 12 control placentae and 14 GDM placentae). Data were analyzed by Mann-Whitney t-tests. *p <0.05, **p< 0.01. Related to Figures 4 and 5.

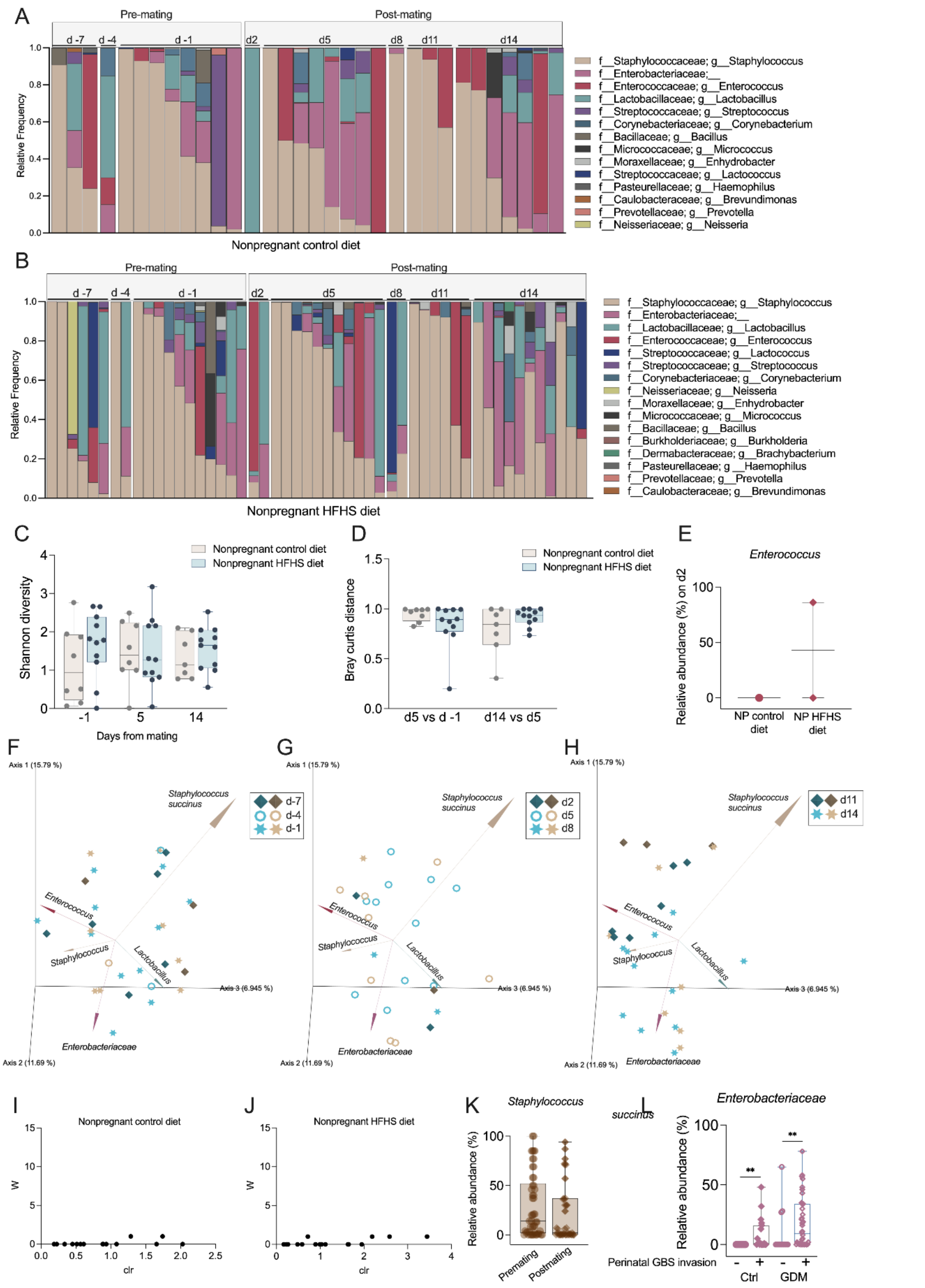


**Figure S5: Murine vaginal microbiota dynamics in non-pregnant cage mates on HFHS and control diets.** A) non-pregnant control-fed and **B)** non-pregnant HFHS-fed mice were swabbed prior every three days from one week before mating (d-7) until d14

just prior to GBS challenge, and longitudinal vaginal microbiome composition was determined by 16Sv4 rRNA amplicon sequencing. **C)** Shannon entropy of vaginal communities at d-1, d5, and d14 timepoints. **D)** Bray Curtis distances between early (d-1 and d5) and late (d5 and d14) timepoints paired per mouse for non-pregnant mice, showing no effects of diet over the time course. **E)** *Enterococcus* relative abundance 2 days after mating, showing GDM specific effects compared to data in Fig. 7. **F-H)** Bray Curtis distance matrix principle coordinate analysis (PcoA) plots of vaginal communities in non-pregnant mice at **F)** pre-mating, **G)** early, and **H)** mid-experimental timepoints, with gray symbols representing controls samples and blue symbols representing mice on HFHS diet. Analysis of Composition of Microbes (ANCOM) in non-pregnant mice on **I)** control or **J)** HFHS diet across all timepoints. **K)** Relative abundance of *S. succinus* at pre-mating and post-mating timepoints. **L**) Relative abundances of *Enterobacteriaceae* at early gestation (E2, E5, E8) timepoints in pregnant control and GDM dams, stratified by GBS dissemination showing earlier effects in line with later timepoint data shown in Fig. 7. Points represent individual samples and lines indicate medians. Box and whisker plots extend from $25^{th}$ to $75^{th}$ percentiles and show all points (C,D,K,L). Data are from 4 independent experiments, $n$ = 3-8 non-pregnant mice on control diet per timepoint, $n$ = 6-11 non-pregnant mice on HFHS diet per timepoint. Experimental numbers for pregnant mice are given in Fig. 7. Data were analyzed by Mann-Whitney t-test (C-E, K-L). **p< 0.01.
